# Transcriptional reprogramming by IL-2 variant generates metabolically active stem-like T cells

**DOI:** 10.1101/2023.05.24.541283

**Authors:** Yaquelin Ortiz-Miranda, Maria Masid, Cristina Jiménez-Luna, Galia Magela Montalvo Bereau, Tania Muller, Nicolas Rayroux, Elisabetta Cribioli, Jesús Corría-Osorio, Helen Carrasco Hope, Romain Vuillefroy de Silly, Bili Seijo, Pierpaolo Ginefra, Kalet León, Nicola Vannini, Ping-Chih Ho, Isaac Crespo, Vassily Hatzimanikatis, Melita Irving, George Coukos

## Abstract

Interleukin-2 receptor (IL-2R)-mediated intracellular signaling is a key regulator of T-cell fate decisions. While the potent signals generated by IL-2 engagement execute effector differentiation, elevated metabolic activities and rapid cellular expansion, IL-15 binding induces a stemness/memory phenotype and a quiescent metabolic state. Here, we demonstrate that weak but sustained signaling generated by a non-IL-2R*α*-binding variant of IL-2 (IL-2v) drive proliferation/metabolic and stemness transcriptional programs, thereby reprogramming CD8^+^ T cells into a hybrid ‘metabolically active stem-like state’. We further show that IL-2v-induced T cells are capable of superior engraftment, persistence, and tumor control when utilized in adoptive cell therapy. Taken together, our study highlights the ability to fine-tune cytokine engagement of cognate receptors in order to generate therapeutically relevant T-cell states and further reveals the metabolic plasticity of the T-cell memory program.

## Introduction

The IL-2R can assemble in two distinct configurations and serves as a rheostat for T-cell activation and differentiation upon ligand engagement. In its heterodimeric format on naïve and memory T cells, the IL-2R*β* (CD122) and the common gamma chain (*γ*_c_ /CD132) engage IL-2 or IL-15 with similar intermediate-affinity (Kd ∼10^−9^M). Upon T-cell activation, the non-signaling IL-2R*α* chain (CD25) is upregulated, binds IL-2 with low-affinity (Kd ∼10^−8^M), and subsequently triggers the formation of a high-affinity heterotrimeric complex (Kd ∼10^−11^M) ^1–3^. Assembly of the trimeric IL-2R*αβγ* complex by IL-2 and consequent signaling promotes clonal T-cell expansion, effector differentiation, and a metabolically active cell state. In contrast, ligand engagement of IL-2R*βγ* favors memory/stemness, limited expansion, and a metabolically quiescent cell state ^4^.

Adoptive T-cell Therapy (ACT) has been hailed as a breakthrough in cancer immunotherapy, and IL-2R stimulation has been crucial to its success ^5, 6^, both by promoting T-cell expansion *ex vivo* and by supporting T-cell persistence upon co-administration *in vivo* ^7–14^. However, unfavorable properties of IL-2, including terminal differentiation of effector T cells^15–18^, induction of activation-induced cell death (AICD)^9, 19, 20^, expansion of regulatory T (Treg) cells, and toxicity at high doses in patients, have driven the development of IL-2 variants with modified biological properties^21–25^. To date, precisely how IL-2 variants alter cell fate decisions is not well understood, nor is it known whether they induce alternative and more efficient metabolic and transcriptomic T-cell states.

In this study, we optimized IL-2/IL-2R interactions in T cells by substituting IL-2/IL-15 with an engineered IL-2R*βγ* agonist, rationally modified to abrogate IL-2R*α* binding^26, 27^ (henceforth referred to as IL-2 variant; IL-2v). Using a comprehensive systems biology approach including mathematical modeling and high-dimensional analyses of metabolic tasks and gene regulatory network inference, we reveal that IL-2v alters T-cell fate decisions to drive a previously undescribed metabolically active stem-like T cell state (abbreviated T_MAS_) that differs from both natural IL-2- and IL-15-induced cell states. Moreover, we demonstrate that these effects confer superior tumor control to CD8^+^ T cells, either expanded in the presence of IL-2v or gene-modified for enforced secretion of IL-2v. Taken together, this study deepens our molecular and cellular understanding of the effects of IL-2R signaling in T-cell fate decisions and metabolic adaptations. Moreover, it underscores the value of ligand/receptor engineering to alter the biological behavior of T cells for therapeutic purposes.

## Results

### IL-2v couples rapid cell expansion to stemness

During canonical CD8^+^T-cell activation, signals are received and integrated from the T-cell receptor (TCR), CD28 and the IL-2R. However, *in vivo*, T cells are further exposed to a diverse set of cues, which vary depending on the inflammatory context^28^. Such complexity curtails the precise investigation of IL-2 versus IL-2v on the biology of CD8^+^T-cells. To dissect the effect of perturbations in IL-2R mediated signals in an unbiased way, we used murine OT1-TCR CD8^+^T cells specific for the ovalbumin-derived SIINFEKL peptide. T cells were activated with anti-CD3/CD28 beads upon harvest while exposed for up to 10 days to IL-2v vs. wild-type IL-2 (**Figure 1A** and **1B**). We included IL-15 as a control ligand of IL-2R*βγ;* like IL-2v, IL-15 does not engage the IL-2R*α* chain^29^. IL-2v is a weak IL-2R agonist and it is not active in the same dose range as IL-2, nor IL-15. Thus, we compared equipotent concentrations (IU/mL) of the cytokines as determined by an IL-2/IL-15 gold-standard fluorometric proliferation assay (**See Methods**, **Figure S1A-S1C**)^26^.

**Figure 1.**
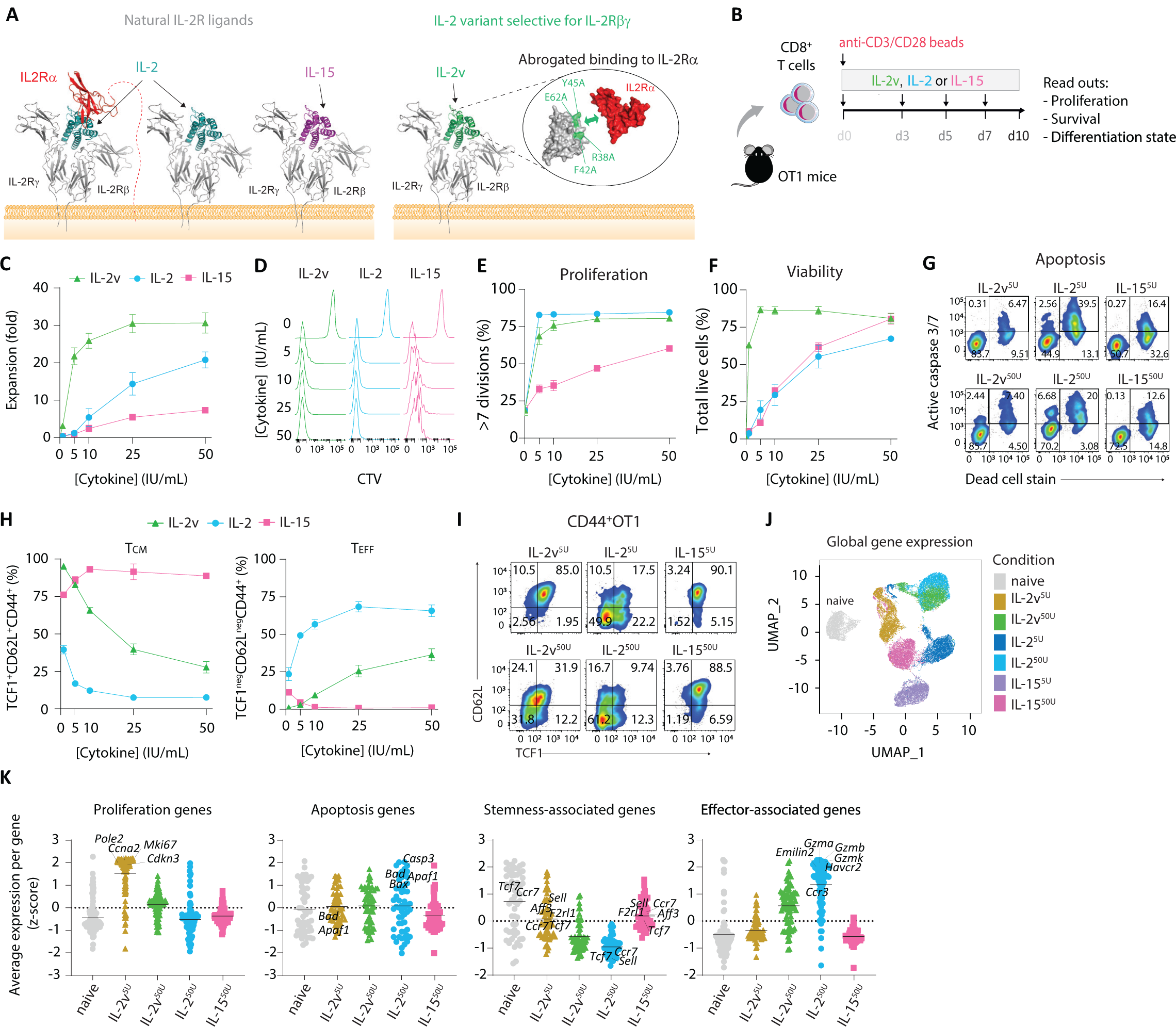
IL-2v couples CD8^+^T cell proliferation/survival to memory differentiation. (**A**) IL-2R complex with natural ligands and IL-2v. (**B**) Schematic of activation and expansion of naïve splenic OT1-T cells. Dose response effects of IL-2v, IL-2 and IL-15 on **(C)** fold-expansion, **(D)** division (CTV dilution), **(E)** proportion of cells undergoing greater than 7 divisions, and **(F)** viability, on day 7. (**G**) Representative dot plots of apoptotic T cells. **(H)** Dose-response effect of IL-2v, IL-2 and IL-15 on the frequency of TCM and TEFF OT1-T cells on day 7, and (**I**) representative dot plots within CD8^+^CD44^+^ T cells. (**J** and **K**) scRNA-seq analysis of naïve and 10-day *in vitro* expanded (as in **(B)**) OT1-T cells. (**J**) UMAP plot showing the transcriptional profiles of OT1-T cells based on global gene expression. (**K**) Expression of proliferation (metaPCNA signature), apoptosis (KEGG apoptosis signature), stemness-associated (UP in LCMV Armstrong d30 vs d8), and effector-associated (DOWN in LCMV Armstrong d30 vs d8) genes in OT1-T cells (refer to Extended Data, Table 1 for gene signatures). Data show mean ±SEM. For (**c-i**), n=3 biological replicates, representative of 3-4 independent experiments. TCM, central memory; TEM, effector memory; TEFF, effector; CTV, CellTrace Violet; scRNA-seq, single cell RNA sequencing; UMAP, Uniform Manifold Approximation and Projection. Refer to Figure S1 for additional information.

We first investigated IL-2v’s ability to promote CD8^+^ T-cell expansion *in vitro* (**Figure 1B**). Strikingly, IL-2v at equipotent concentrations induced 2-4-fold higher OT1 T-cell expansion than IL-2, and 5-6-fold higher expansion than IL-15 (**Figure 1C**). The low expansion by IL-15 is consistent with its role in driving quiescence *in vivo*^30^. Importantly, IL-2v and IL-2 induced similar levels of T-cell proliferation (**Figure 1D** and **1E**), but T-cell survival/viability was higher with IL-2v (**Figure 1F** and **1G**). Thus, IL-2v produces distinct biological outcomes from IL-2 and IL-15.

While IL-2 is known to drive both effector differentiation and rapid T-cell expansion^4^, we sought to determine whether IL-2v would instead promote stemness like IL-15^30^. Indeed, IL-2v increased the proportion of central memory (T_CM_) cells relative to IL-2 (**Figure 1H, 1I** and **S1D**), which instead favored effector (T_EFF_) differentiation as expected^31, 32^. Strikingly, IL-2v applied at a low concentration (5 IU/mL; hereafter referred to as IL-2v^5U^) promoted comparable overall T-cell expansion relative to 50 U/mL of IL-2 (IL-2^50U^) and induced similar memory differentiation as IL-15, with more than 80% of cells being T_CM_ after expansion (**Figure 1C**). Thus, unlike IL-2 which drives high proliferation coupled with effector differentiation, or IL-15 which favors memory differentiation and quiescence, IL-2v^5U^ acts as a hybrid, promoting high proliferation along with memory differentiation and enhanced survival, thus coupling T-cell expansion to memory differentiation.

Single-cell (sc) RNA-seq of OT1-T cells cultured for 10 days with 5 or 50 IU/mL of IL-2v, IL-2 or IL-15 concorded with the above results. IL-2v^5U^-OT1-T cells clustered independently from IL-2- or IL-15-, and IL-2v^50U^-OT1-T cells (**Figure 1J**) and were characterized by high expression of proliferation-associated genes (e.g., *Mki67, Cdkn3* and *Ccna2*) as well as memory/stemness-associated genes (e.g., *Sell, Tcf7* and *Ccr7*) (**Figure 1K** and **Table S1**). Unlike IL-2^50U^, IL-2v^5U^-OT1-T cells showed low expression of genes involved in cell migration (e.g.*, Emilin2, Lgals3* and *Ccr3Mi*), effector function (e.g.*, Gzmb, Gzma* and *Ccr3*), and apoptosis (e.g.*, Tnfrsf10b, Bad* and *Casp3*) (**Figure 1K** and **Table S1**). Overall, IL-2v^50U^ induced an intermediate state compared to IL-2v^5U^ and IL-2^50U^, driving higher memory differentiation than IL-2^50U^ while partially upregulating effector genes in contrast to IL-2v^5U^.

In summary, IL-2v instructs the acquisition of transcriptional states that differ from those induced by IL-2 or IL-15 in a dose-dependent manner. We conclude that IL-2v^5U^ induces a stem-like T-cell state characterized by high cell proliferation and survival while simultaneously retaining stemness.

### IL-2v generates metabolically active memory T cells

We next sought to explore the metabolic processes underpinning the stem-like T-cell state induced by IL-2v^5U^ that could explain the rapid cell expansion. UMAP representation of all metabolic genes from the above scRNA-seq data separated IL-2v^5U^ from canonical IL-2 or IL-15 (**Figure 2B**). In comparison, IL-2v^5U^ upregulated the expression of enzymes from the citric acid cycle (TCA), oxidative phosphorylation (OXPHOS), the pentose phosphate pathway (PPP), and the nucleotide synthesis pathways, while the expression of glycolytic and fatty acid oxidation (FAO) enzymes were downregulated (**Figure S2B**). Moreover, IL-2v^5U^ upregulated a distinct set of cell membrane and mitochondrial transporters relative to IL-2^50U^ or IL-15^50U^ (**Figure S2C and S2D**). These results suggest IL-2v induced a different metabolic state than IL-2 and IL-15.

**Figure 2.**
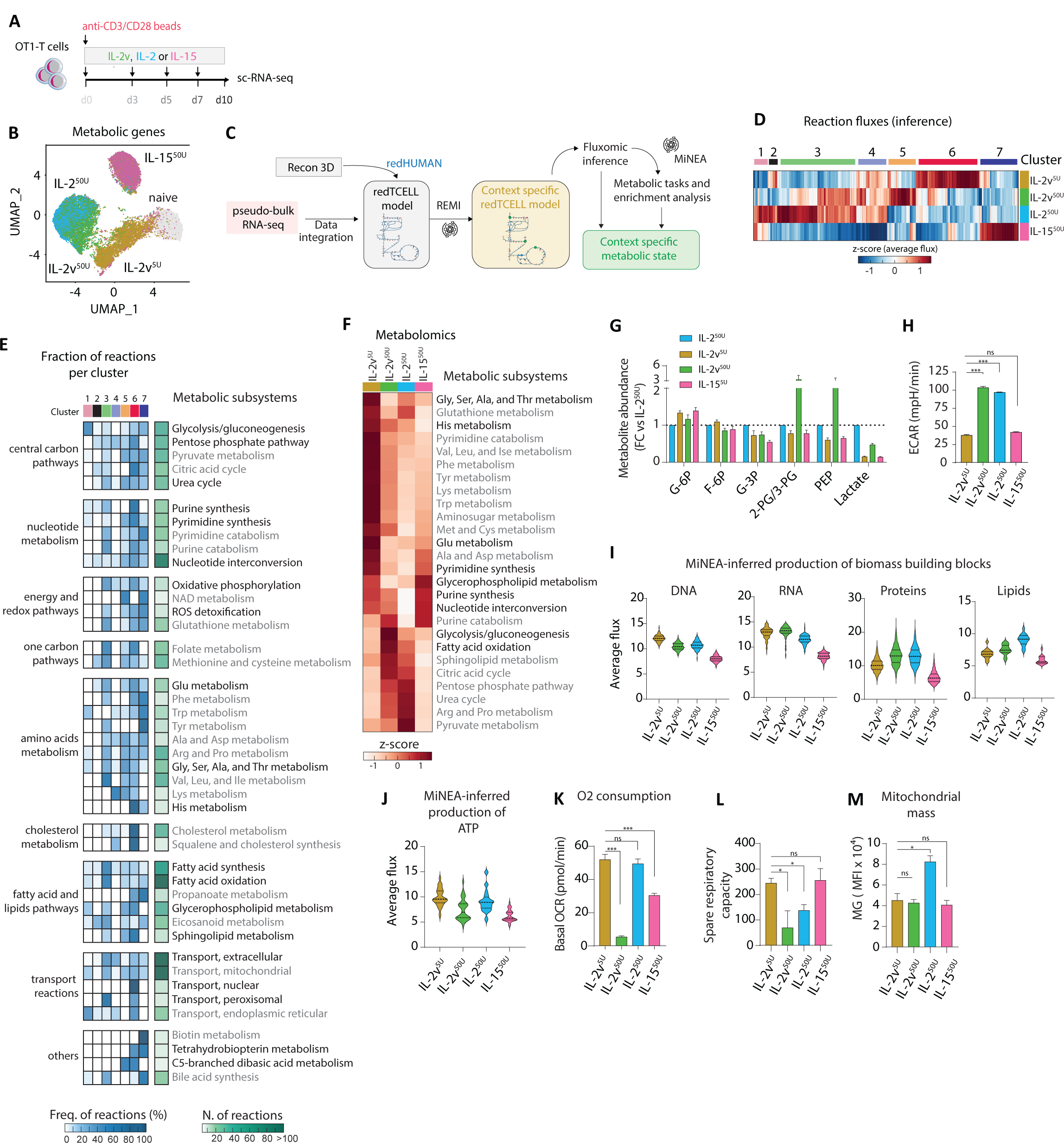
IL-2v generates metabolically active stem-like T cells without upregulating aerobic glycolysis and while maintaining a high level of mitochondrial fitness. (**A**) Schematic of naïve OT1-T cell activation and stimulation prior to functional measurements on day 10 of expansion. (**B**) UMAP plot showing transcriptional profiles based on metabolic gene expression. (**C**) Schematic of *in silico* fluxomics pipeline. (**D**) Heatmap showing average flux values for all metabolic reactions present in the redTCELL metabolic model (See Methods for details). (**E**) Heatmap showing the fraction of reactions pertaining to each metabolic subsystem included in the clusters shown in (**D**) (left, in blue) and total number of reactions per metabolic subsystem (right, in green). (**F**) Heatmap showing enrichment of metabolic pathways based on the relative abundance of 167 central carbon metabolites in OT1-T cells. (**G**) Fold change of the abundance of glycolytic intermediates relative to the IL-2^50U^ condition. (**H**) Representative ECAR measured by Seahorse in real time under basal conditions. (**I** and **J**) Inferred production of (**I**) DNA, RNA, proteins, lipids, and (**J**) ATP (by the mitochondrial ATP synthase) based on MiNEA (refer to Methods for details). (**K** and **L**) Representative OCR measured by Seahorse in real time under (**K**) basal conditions only or (**L**) also in response to the mitochondrial inhibitors oligomycin, FCCP, and R/A. (**L**) Spare respiratory capacity. (**M**) Mitochondrial mass based on MG staining. (**N**) Relative regulation of metabolic genes pertaining to the indicated metabolic subsystems by proliferation- and effector-associated TFs (depicted in Figure 3F and **3H**). Data show mean ±SEM. For (**F** and **H**), n=4 biological replicates. For (**I**, and **K-N**), n=3 biological replicates, representative of 2-3 independent experiments. *p<0,05, **p<0,01 ***p<0,001, ****p<0,0001 using Unpaired t test with Welch’s correction. UMAP, Uniform Manifold Approximation and Projection; MiNEA, minimal network enrichment analysis; FC, fold change; OCR, O2 consumption rate; FCCP, carbonyl cyanide 4-(trifluoromethoxy) phenylhydrazone; ECAR, extracellular acidification rates; R/A, rotenone and antimycin; MG, MitoTracker green. Refer to Figure S2 for additional information.

To further understand the IL-2v^5U^ novel metabolic state at a systems level, we conducted *in silico* fluxomic analysis to infer from gene expression information the rate of metabolic reactions (fluxes) induced by each cytokine condition, which was validated by intracellular pan-metabolite analysis by MS. We first applied redHUMAN^33^ to construct a T-cell-adapted metabolic model (redTCELL) comprising biochemical and thermodynamic information of core metabolic subsystems (**Figure 2C**, **See Methods** and **Table S3**). We integrated our scRNA-seq data in redTCELL to infer the flux for each metabolic reaction per condition (**Figure 2C**). Relative to IL-2^50U^, IL-2v^5U^ upregulated TCA, PPP, nucleotide synthesis, structural lipids metabolism (e.g., cholesterol, glycerophospholipids and sphingolipids), amino acid metabolism and nuclear transport fluxes (**Figure 2D, 2E** and **S2F**). Notably, IL-2v^5U^ upregulated the majority of metabolic fluxes relative to IL-15^50U^, indicating a more active metabolic state despite the maintenance of stemness characteristics (**Figure 2D and 2E**). Enrichment analysis based on MS-measured metabolites corroborated these inferences (**Figure 2F**). However, IL-2v^5U^ downregulated the rates of aerobic glycolysis, FAO, fatty acid synthesis (FAS) and peroxisomal transport fluxes as compared to IL-2 and IL-2v^50U^ (**Figure 2D, 2E** and **S2F**). Also consistent with lower activation of aerobic glycolysis by IL-2v^5U^, we found lower abundance of glycolytic intermediates (**Figure 2G**) and lower extracellular acidification rate (ECAR) in IL-2v^5U^-relative to IL-2^50U^-T cells (**Figure 2H**).

Because central carbon pathways such as glycolysis, TCA and PPP contribute to anabolic pathways, redox balance, and energy production, we thought to infer the relative synthesis of cell growth macromolecules (RNA, DNA, proteins, and lipids) given the differential regulation of fluxes induced by IL-2v^5U^ relative to IL-2^50U^ and IL-15^50U^. Thus, we used redTCELL to identify the critical metabolic reactions required to fulfill these metabolic tasks and performed enrichment analysis using the above *in silico* fluxomic data (**Figure 2C** and **See Methods**)^34^. IL-2v^5U^ resulted in a greater capacity to generate growth macromolecules in T cells than IL-15 (**Figure 2I**) consistent with the variant’s enhanced capacity to enhance CD8^+^T-cell proliferation (**Figure 1C** and **1E**). Moreover, IL-2v^5U^ induced higher flux through reactions required for the synthesis of DNA and RNA than IL-2^50U^, but lower flux for the synthesis of protein and lipids (**Figure 2I**), explaining the 2-fold smaller cell size under IL-2v^5U^ expansion (**Figure S2E**). Moreover, we inferred from the metabolic network that IL-2v^5U^ and IL-2^50U^ induced similar ATP production by the mitochondrial ATP synthase, which was ∼1.5-fold higher than IL-15^50U^ (**Figure 4J**). Consistently, IL-2v^5U^-T cells exhibited comparable baseline oxygen consumption rate (OCR) to IL-2^50U^-T cells (**Figure 2K**), while IL-15^50U^-T cells yielded lower values. Interestingly, IL-2v^5U^, like IL-15^50U^-T cells, retained high spare respiratory capacity (SRC) (**Figure 2L**), and a smaller mitochondrial mass than IL-2^50U^-T cells (**Figure 2M**), suggesting that IL-2v^5U^ confers high mitochondrial fitness despite the enhanced metabolic activity.

**Figure 4.**
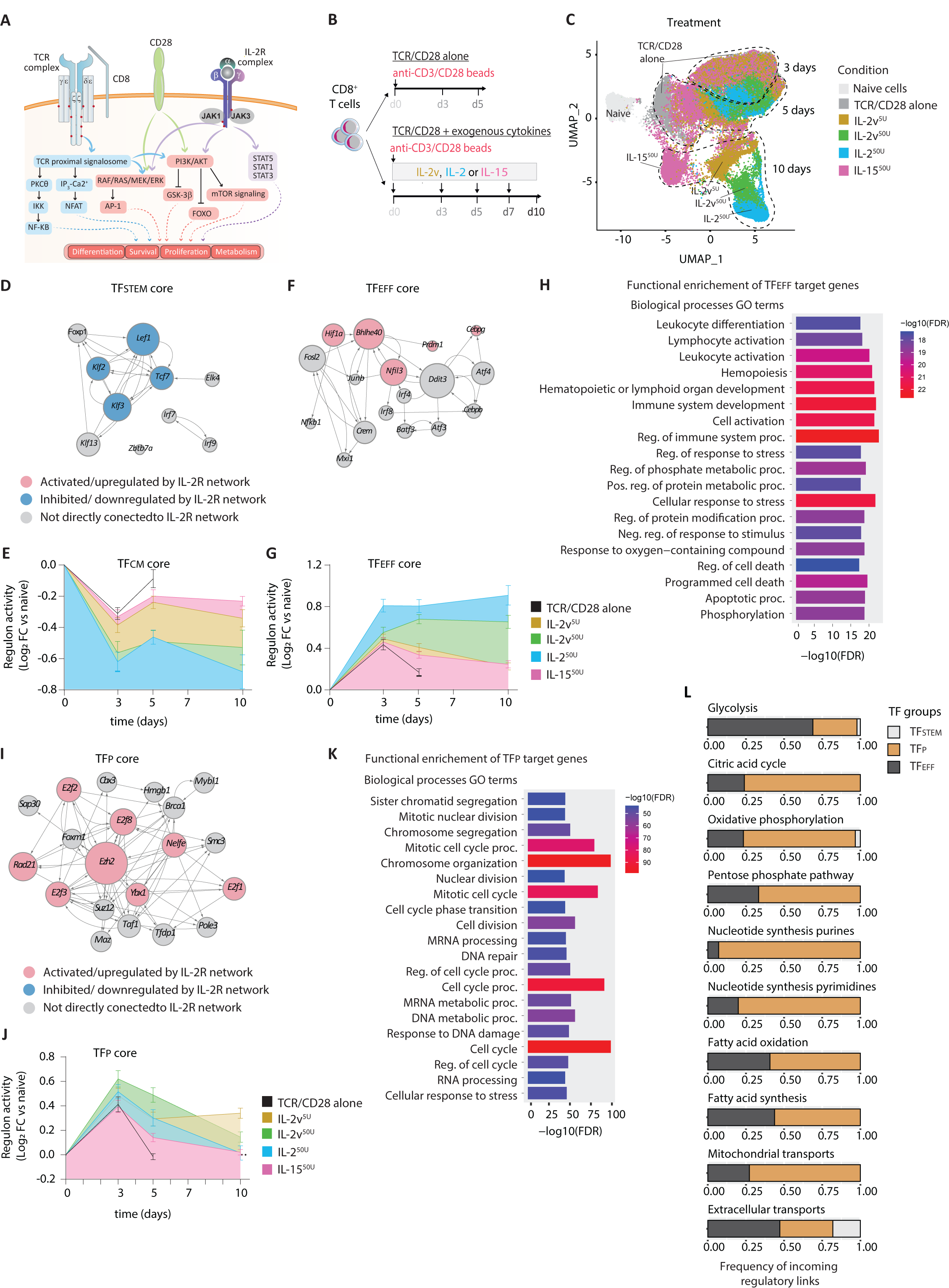
IL-2v co-activates stemness and proliferation transcriptional programs. **(A)** Schematic of IL-2R crosstalk with TCR and CD28 signaling pathways. **(B)** Schematic of naïve OT1-T cells activation and stimulation during expansion for sample collection on days 0, 3, 5 and 10 to perform scRNA-seq. **(C)** UMAP plot showing the transcriptional profiles of OT1-T cells based on global gene expression. (**D, F** and **I**) Subnetworks of (**D**) stemness(TFSTEM)-, (**F**) effector(TFEFF)- and (**H**) proliferation(TFP)-associated regulons (refer to Methods for details). Node size represents out degree in the global gene regulatory network. (**E, G** and **J**) Kinetics of regulon activity per group of (**E**) stemness(TFSTEM)-, (**G**) effector(TFEFF)- and (**I**) proliferation(TFP)-associated regulons. (**H** and **K**) GO functional enrichment analysis of target genes from (**H**) TFEFF or (**K**) TFP. (**L**) Frequency of regulatory links between the indicated TF groups and genes pertaining to the indicated metabolic subsystems. scRNA-seq, single cell RNA sequencing; UMAP, Uniform Manifold Approximation and Projection; TF, transcription factor; FC, fold change; AUC, activity unit per cell; and GO, gene ontology. Refer to Figure S4 and S5 for additional information.

Taken together, we have shown that IL-2v^5U^ generates stem-like T cells that are endowed with the ability to generate the biomass and energy needed to sustain rapid cell expansion without upregulating aerobic glycolysis. Hereafter we refer them as ‘metabolically active stem-like T cells’ (T_MAS_). In contrast, IL-2^50U^ drives effector differentiation and upregulates glycolysis while IL-15^50U^ generates conventional memory T cells having a quiescent metabolic state.

### Low and persistent IL-2R signaling by IL-2v generates T_MAS_ cells

Because IL-2v and IL-2/IL-15 utilize the same IL-2R signaling dimer (IL-2R*βγ*_c4, 35_), we hypothesized that dissimilarities in T-cell fate decisions were due to qualitative or/and quantitative differences in signaling. The signaling networks downstream of the IL-2R include Janus kinase (JAK)-signal transducer and activator of transcription (STAT), mitogen-activated protein kinase (MAPK), and the phosphoinositide 3-kinase (PI3K)-protein kinase-B (AKT) pathways^4^ (**Figure S3A** and **S3G**). We investigated the activation levels of these pathways in OT1-T cells following a 20-minute cytokine pulse. IL-2v induced lower phosphorylation of JAK1 and JAK3 (most upstream mediators) (**Figure S3B-S3D**), as well as of STAT5, ERK1/2, and AKT (major hubs for proximal IL-2R signaling cascades) (**Figure 3B** and **3C**). These were however similar to or higher than for IL-15, depending on dose (**Figure 3B** and **3C**). We observed similar results with human CD8^+^T cells, indicating strong conservation of these effects (**Figure S3E** and **S3F**). To explore signaling in a more comprehensive and unbiased manner, we carried out a phospho-proteome wide analysis by mass spectrometry (MS) under the same 20-min cytokine pulse conditions and applied enrichment analysis to identify relative abundances of 390 identified phospho-species known to partake in IL-2R pathways. Relative to unstimulated cells, IL-2v triggered lower overall activation of IL-2R-associated signaling networks than IL-2, and this was more pronounced for IL-2v^5U^ (**See Methods**, **Figure 3D, S3G** and **S3H**).

**Figure 3.**
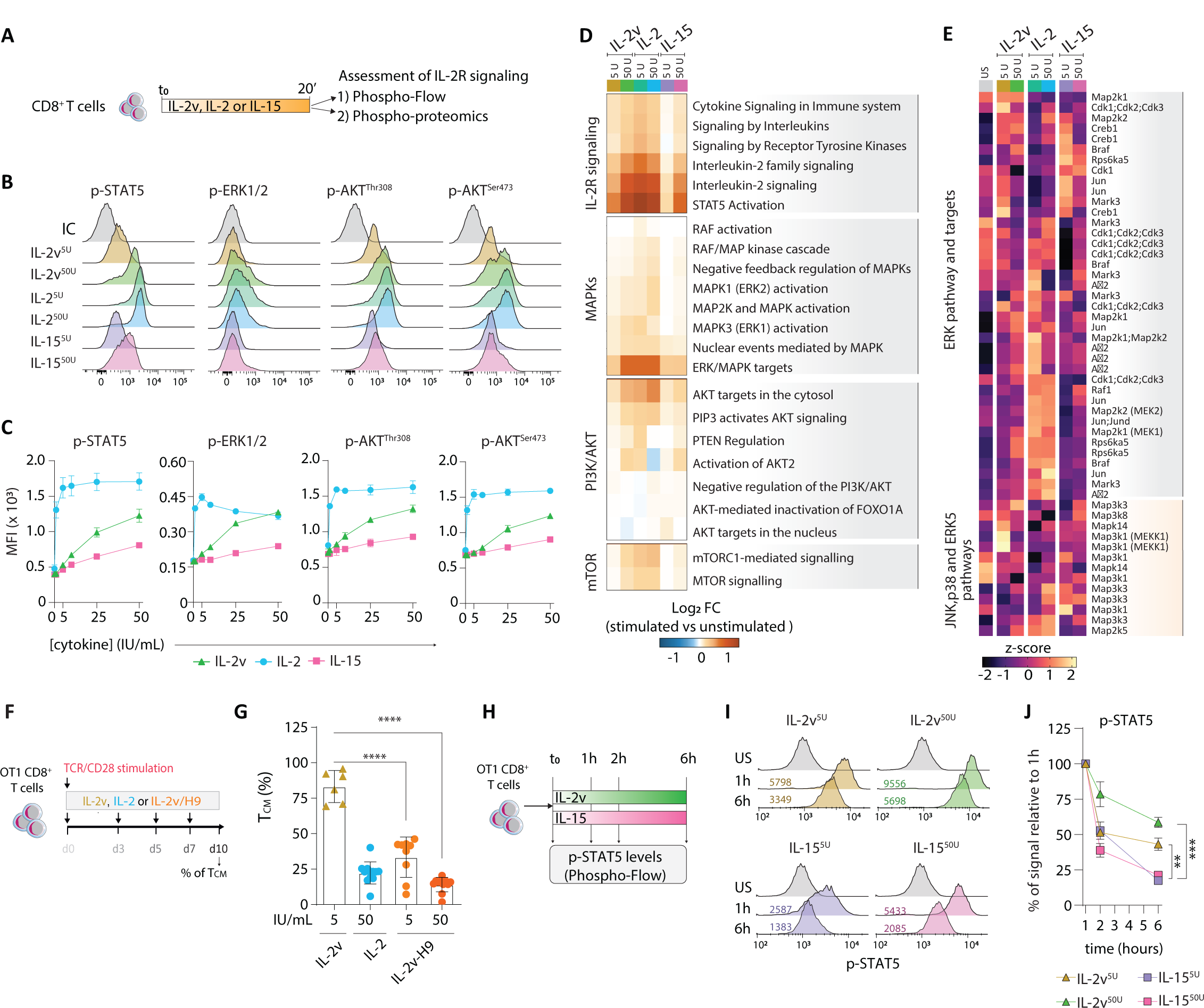
IL-2v induces a lower-intensity tonic activation of IL-2R signaling networks compared to wild-type IL-2. (**A**) Schematic of OT1-T cell cytokine stimulation prior to phospho-flow cytometry (**B** and **C**) or phospho-proteomics (**D** and **E**). (**B**) Representative histograms and (**c**) dose-response effect of cytokines on the phosphorylation of signaling molecules. (**D**) Activity of IL-2R proximal pathways based on phospho-site enrichment analysis. (**e**) Phosphorylation levels MAPK species. (**F**) Schematic of naïve OT1-T cell stimulation and expansion. (**G**) Frequency of OT1-TCM cells (cultured as in **(F)**) at day 10 of expansion. (**H**) Schematic of OT1-T cell stimulation prior to phospho-flow cytometric analysis (**I** and **J**). (**I**) Representative histograms and (**J**) cumulative data of p-STAT5 kinetics post-stimulation. Data show mean ±SEM. For (**C** and **J**), n=3 biological replicates; representative of 3 independent experiments. For (**D** and **E**), n=3 biological replicates. For (**G**) n=6-9 biological replicates; cumulative of 3 independent experiments. *p<0,05, **p<0,01 ***p<0,001, ****p<0,0001 using Unpaired t test with Welch’s correction. Abbreviations: TCM, central memory; IL-2v-H9, IL2 variant with IL-2v and H9-superkine mutations. Refer to Figure S3 for additional information.

Activation of the AKT network is critical for effector differentiation and metabolic adaptations in T cells, including aerobic glycolysis and protein synthesis to support effector functions. Consistent with lower AKT phosphorylation, IL-2v induced lower activation of mTORC1 signaling (a key downstream mediator controlling protein synthesis and glycolysis), explaining the lower glycolytic rate in IL-2v^5U^-relative to IL-250U-OT1-T cells (**Figure 2E-2H**). Besides, IL-2v induced lower phosphorylation of cytosolic or nuclear AKT targets, including FOXO1/3a and GSK-3*β*, than IL-2 (**Figure 3D** and **S3I-S3K)**. Moreover, IL-2v induced lower phosphorylation levels of pro-apoptotic proteins in OT1-T cells than IL-2 (**Figure S3H**). Intriguingly, based on phospho-site abundance, the lower enrichment of MAPK subnetworks was not the result of a global decrease in phosphorylation of all intermediaries. Indeed, we observed higher activation of certain species by IL-2v, including cyclin-dependent kinases 1, 2 and 3 (Cdk1-3) relative to IL-2 (**Figure 3E**). Overall, IL-2v^5U^ stimulation resulted in attenuated IL-2R signaling as compared to IL-2.

To validate if IL-2v driven T-cell stemness is dependent on signaling strength, we introduced on the IL-2R*β* binding domain of IL-2v four amino acid replacements from the well-characterized H9 superkine, which has a 200-fold higher binding affinity for IL-2R*β*^36^. The IL-2v/H9 variant, expected to engage IL-2R*βγ* with high affinity but not IL-2R*α* (**Figure S3L**), exhibited similar activity to H9 in a fluorometric proliferation assay using CTLL-2 cells, indicating similar IL-2R binding strength (**Figure S3M**). Importantly, IL-2v/H9 variant exhibited higher overall signaling intensity than IL-2v in OT1-T cells (**Figure S3N**), which resulted in the loss of memory differentiation (stemness) (**Figure 3F** and **3G**). Thus, low-intensity IL-2R signaling, conferred to IL-2v by its inability to bind IL-2R*α*, is key for inducing stemness.

Finally, we sought to understand how IL-2v^5U^ generates metabolically active, highly proliferative CD8^+^ T-cells relative to IL-15^50U^ despite a similar signaling strength and memory differentiation. We theorized that the difference could be due to signal duration. To address this question, we analyzed STAT5 phosphorylation dynamics following stimulation with IL-2v or IL-15 at equipotent doses (5 and 50 IU/mL). Notably, STAT5 phosphorylation persisted at higher levels 6 hours following exposure to IL-2v than IL-15 (**Figure 3H-3J**). Thus, IL-2v^5U^ induces low-intensity, persistent (tonic/durable) signals in CD8^+^T cells, promoting the novel metabolically active stem-like T cell state allowing high proliferation. In contrast, the strong signaling generated with IL-2 promotes metabolically active effector cells, and the low-intensity, short-lived signals from IL-15 yields metabolically quiescent memory T cells.

### IL-2v sustains transcriptional programs governing stemness and proliferation/metabolic activity

Next, we sought to elucidate how signaling mediated by IL-2v affects transcriptional regulation of CD8^+^T-cell fate decisions to enable the generation T_MAS_ cells. IL-2R cross-talks with TCR and CD28 signaling networks at the MAPKs and PI3K/AKT levels (**Figure 4A**). We thus hypothesized that different intensity IL-2R signals fine-tune the signaling triggered by TCR/CD28 receptors, instructing transcriptional regulation depending on signal strength and duration. We analyzed by scRNA-seq OT1-T cells expanded with IL-2v^5U^ or IL-2v^50U^ early during activation (3 days) and after TCR/CD28 signals had ceased (5 and 10 days) (**Figure 4B, S4A** and **S4B**). As a control we analyzed OT1-T cells primed with TCR/CD28 but cultured without cytokine stimulation. Although the cell states were similar on day 3, these separated as the TCR/CD28 effects faded, revealing a distinct dose-dependent transcriptional state driven by IL-2v, which also differed from those induced by IL-2 or IL-15 (**Figure 4C**).

To unveil the activity of transcription factors (TFs) instructing fate decisions, we used the scRNA-seq data to construct a global gene regulatory network, from which we inferred 819 TF regulons (units of TF and target genes) (**See methods**). First, we focused on fate decisions associated with memory (stemness) versus effector differentiation. To that end, we identified TFs linked to either of these alternative fates considering expression levels and regulon activity. TFs with the highest discriminatory power for T_CM_ were defined as stemness-associated TFs (TF_STEM_) and for T_EFF_ cells as effector-associated TFs (TF_EFF_) (**See methods** and **Figure S4C-S4E**). Seven of the ten identified TF_STEM_ formed a regulatory core (**Figure 4D**) and seventeen of the twenty identified TF_EFF_ regulons were also cross-regulated, forming another regulatory core (**Figure 4F**), implying that activation of some of these nodes could propagate through these networks, activating the set of regulons. Four central regulons to the TF_STEM_ core, *Klf2*, *Tcf7*, *Lef1* and *Klf3* (**Figure 4D**) are critical regulators of stemness programs in memory and naive cells^37^, and are negatively regulated by AKT and STAT5 signaling^38, 39^. In contrast, within the TF_EFF_ core, *Hif1a*, *Bhlhe40*, *Nfil3*, *Prdm1*, and *Cbpg* (**Figure 4F**) are known for their roles in effector commitment and are positively regulated by AKT/mTORC1 and STAT5^38, 40^.

Functional annotation of target genes of these TF_EFF_ revealed that they indeed regulate genes involved in lymphocyte activation, differentiation, and response to stress/stimulus (**Figure 4H**). We thus reasoned that only strong IL-2R signals inducing powerful activation of the AKT/mTORC1 and STAT5 signaling networks, provided by IL-2 but not IL-2v^5U^, would lead to stable downregulation of the TF_STEM_ core and upregulation the TF_EFF_ core. Indeed, in contrast to IL-2^50U^, and like IL-15^50U^, low-intensity IL-2v^5U^ had no effect on TCR/CD28-driven TF_STEM_ inhibition or TF_EFF_ activation during priming and favored TF_STEM_ over TF_EFF_ activation at later time points (**Figure 4E, 4G, S4G and S4H**), hence explaining the memory commitment upon TCR/CD28 signal fading. Interestingly, IL-2v^50U^ signals inhibited TF_STEM_ and activated TF_EFF_, but to a lesser extent than IL-2^50U^, emphasizing the role of signal strength in IL-2v’s effects and correlating with the intermediate levels of stemness induced by this dose of the variant. Thus, IL-2R signal strength appears to determine the preferential activation of memory vs. effector transcriptional programs.

To corroborate our interpretation that IL-2R signaling strength determines memory/stemness vs. effector transcriptional programs, we activated and expanded OT1-T cells in the presence of IL-2^50U^ and used escalating concentrations of tofacitinib, a JAK inhibitor (JAKi), to progressively attenuate IL-2R downstream signaling (**Figure S5A**). JAKi resulted in a dose-dependent decline in effector differentiation after a week in culture in IL-2^50U^ (**Figure S5B and S5C)**. Importantly, we observed a similar decline in the mRNA expression levels of the above inferred TF_EFF_, already on day 3 (**Figure S5E**). Thus, IL-2R signaling strength regulates TF_EFF_ expression levels and their activation is triggered by strong IL-2R signals provided by canonical IL-2 but not IL-2v^5U^.

To identify the transcriptional program underlying IL-2v^5U^’s ability to promote rapid CD8^+^T cell expansion, we inferred proliferation-associated TFs (TF_P_) based on their correlation with the expression of proliferation-associated genes^41^ in our data set (**Figure S4F** and **See Methods**). TF_P_ formed a regulatory core including well-known TFs or transcriptional regulators (**Figure 4I**). Several TF_P_ are known to be positively regulated by MAPKs and AKT networks downstream of TCR, CD28 and the IL-2R (**Figure 4I**)^42, 43^ and functional annotation of their target genes revealed that they indeed regulate genes involved in cell cycle progression, chromosome segregation, and RNA and DNA metabolic processes (**Figure 4K**). Consistently, cells primed with CD3/CD28 beads without cytokines exhibited upregulation of these TF_P_ after activation, followed by decay within 5 days, commensurate with the decrease in TCR/CD28 signaling (**Figure 4J** and **S4B**). Importantly, IL-2v^5U/50U^ and IL-2^50U^ maintained TF_P_ activation after 5 days (**Figure S4I**) while IL-15^50U^ did not, consistent with the proliferation observed with each cytokine. Strikingly, only IL-2v^5U^ sustained high-level TF_P_ activation at 10 days. In contrast, although IL-2^50U^ signaling triggered TF_P_, it appears that such strong IL-2R signal intensity is not required nor optimal to sustain high proliferation and survival. Notably, strong IL-2^50U^ signals resulted in greater activation of negative regulation of PI3K/AKT and MAPKs pathways than IL-2v (**Figure 3D**) which could result in a long-term dampening of the signal. Strikingly, decreasing IL-2R signal intensity with tofacitinib (JAKi) in OT1-T cells expanded in presence of IL-2^50U^ had no impact on the expression of TF_P_ at 72 hours and was rather beneficial for their expression 48 hours later (**Figure S5A, S5F and S5G**). Altogether, these findings indicate that low-intensity tonic signaling such as provided by IL-2v^5U^ (but not IL-2^50U^ or IL-15^50U^) stabilizes the TF_P_ transcriptional program and sustains proliferation and high cell survival.

Further annotation of the TF_EFF_ and TF_P_ regulons revealed that they regulate not only effector and cell cycle progression genes, but also genes involved in essential metabolic pathways. Conversely, T_STEM_ regulons did not regulate metabolic genes (**Figure 4L**). Importantly, we discovered that TF_P_ regulons preferentially regulate TCA, OXPHOS, and nucleotide synthesis genes compared to TF_EFF_ regulons, which preferentially regulate glycolysis genes (**Figure 4L**). Thus, the selective activation of TF_P_ by IL-2v^5U^ without the upregulation of TF_EFF_ is consistent with the metabolic dysregulation induced by IL-2v^5U^; i.e., the uncoupling of biomass and mitochondrial energy production from aerobic glycolysis (**Figure 2**). Moreover, the potent activation of TF_EFF_ by IL-2^50U^ correlates with the elevated levels of aerobic glycolysis driven by IL-2, and the low-level activation of TF_EFF_ and TF_P_ by IL-15^50U^ is in line with the quiescent metabolic state that IL-15 generates (**Figure 2**). Thus, it appears that low-intensity tonic IL-2R signaling by IL-2v is optimal for driving a coordinated transcriptional and metabolic reprogramming to generated T_MAS_ cells.

### IL-2v expansion improves ACT

Large numbers of minimally differentiated antigen-specific T cells have been demonstrated optimal for ACT as they enable higher *in vivo* persistence and superior tumor control^44, 45^. We reasoned that IL-2v^5U^’s ability to enable both stemness and rapid cell expansion would be ideally positioned for ACT. As expected for memory cells, IL-2v^5U^-OT1-T cells expressed no or low levels of granzyme-B or C, perforin, or coinhibitory receptors (e.g., TIM-3 and LAG-3) (**Figure S6A-S6E**), in contrast to IL-2-OT1 T cells. However, upon *in vitro* restimulation with the OT1-TCR cognate peptide (SIINFEKL), IL-2v^5U^-OT1 cells upregulated IFN*γ* and TNF*α* to comparable levels with IL-2^50U^-OT1 cells, indicating their capacity to acquire effector functions upon TCR restimulation (**Figure S6F**). Interestingly, IL-2v^5U^-OT1-T cells upon co-culture with B16.OVA tumor cells expressing the cognate peptide secreted significantly higher amounts of IFN*γ*, TNF*α* and effector chemokines (e.g., RANTES/CCL5) than IL-2^50U^-OT1-T cells, demonstrating higher polyfunctionality (**Figure 5B**). Furthermore, IL-2v^5U^-OT1-T cells secreted higher amounts of IL-2 (**Figure 5C**), which correlated with higher proliferation and a 300-fold increase in T-cell expansion than IL-2^50U^-OT1-T cells in this setting (**Figure S6G and S6H)**. Finally, IL-2v^5U^-OT1-T cells killed efficiently tumor cells in co-culture (**Figure S6I**). Thus, expanding CD8^+^T cells with IL-2v^5U^ increases polyfunctionality in the presence of target cells, and proliferative capacity *in vitro*.

**Figure 5.**
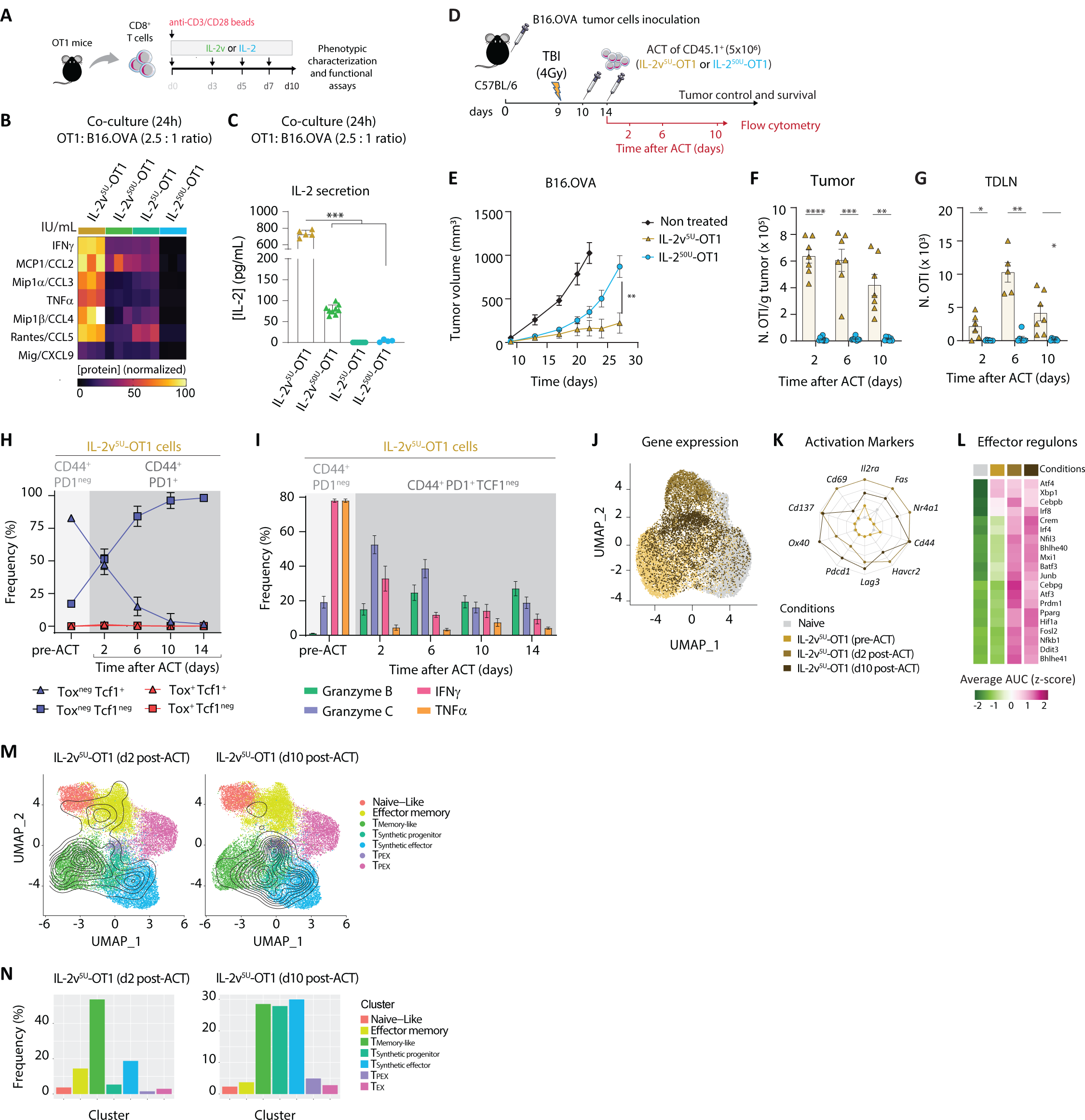
*In vitro* transcriptional reprogramming induced by IL-2v enhances CD8^+^T cell engraftment, persistence, and anti-tumor function. (**A**) Schematic of naïve OT1-T cell activation and stimulation during expansion. (**B** and **C**) Co-culture of OT1-T cells (expanded as in (**A**)) with melanoma B16.OVA tumor cells. (**B**) Heatmap of cytokine and chemokine production, and (**C**) IL-2 production. (**D**) Schematic of ACT *in vivo* study with days of tumor and tissue collection for flow cytometric analysis (**F-I**) or scRNAseq (**J-N**) indicated. (**E**) Tumor growth curves for ACT study. Numbers of persisting OT1-T cells in the (**F**) tumor, and (**G**) tumor draining lymph node post-ACT. (**H**) Cumulative frequencies of TEX (TCF-1^neg^TOX^+^), TPEX (TCF-1^+^TOX^+^), Tprogenitor-like (TCF-1^+^TOX^neg^) and TEFF-like (TCF-1^neg^TOX^neg^) within CD44^+^PD1^+^OT1-T cells in the tumor post-ACT. (**I**) Cumulative frequencies of CD44^+^PD1^+^TCF1^neg^OT1-T cells producing granzyme B and C, IFN*γ* and TNF*α* upon 4-hour re-stimulation with anti-mouse CD3 antibody post-ACT. For (**H** and **I**), pre-ACT (day 10 of *in vitro* expansion) IL-2v^5U^-OT1 (CD44^+^PD1^neg^) cells were used as controls. (**J**) UMAP plot showing transcriptional profiles of IL-2v^5U^-OT1 cells pre-(day 10 of *in vitro* expansion) and post-ACT (days 2 and 10 after second ACT infusion) based on global gene expression. (**K**) Spider plot depicting expression levels of activation markers and (**L**) activity of effector-associated TF regulons for conditions shown in (**J**). (**M** and **N**) Projection of IL-2v^5U^-OT1 cells post-ACT onto OT1/Endogenous transcriptomic space using ProjecTILs (refer to Methods). (**M**) Contour plots depicting the clusters covered by each dataset and (**N**) bar plots showing the cluster composition. Data show mean ±SEM. For (**B** and **C**), n=6-9 biological replicates; pooled data from 3 independent experiments. For (**E**), n=8-10 biological replicates, representative of 3-4 independent experiments. For (**F-I**), n=8-15 biological replicates, pooled data from 3-4 independent experiments. *p<0,05, **p<0,01 ***p<0,001, ****p<0,0001 using Unpaired t test with Welch’s correction. scRNA-seq, single cell RNA sequencing; UMAP, Uniform Manifold Approximation and Projection; ACT, adoptive T cell therapy; TEX, terminally exhausted; TPEX, precursor exhausted. Refer to Figure S6 for additional information.

To assess the ability of T cells expanded under IL-2v^5U^ vs. canonical IL-2 to engraft *in vivo*, we transferred 2×10^6^ OT1-T cells expanded for 10 days in the respective cytokine condition to sublethally irradiated (lymphodepleted) naïve mice (**Figure S6J**). After 24 hours, we observed a 50-fold higher accumulation of IL-2v^5U^-OT1 than IL-2-OT1-T cells in the spleen (**Figure S6K**), with 3-fold more central memory (TCM)-like IL-2v^5U^-OT1-T cells (**Figure S6L**).

We subsequently conducted ACT in sublethally irradiated mice bearing subcutaneous B16.OVA tumors. OT1-T cells were infused twice for a total of 5×10^6^ cells (**Figure 5D**). ACT with IL-2v^5U^-OT1-T cells proved significantly more effective, inducing better tumor control than with IL-2^50U^-OT1-T cells (**Figure 5E**). This effect correlated with enhanced engraftment and persistence of IL-2v^5U^-OT1-T cells in both the tumor and tumor-draining lymph nodes, with >100-fold increase in T-cell accumulation relative to IL-2^50U^-T cells (**Figure 5F** and **5G**).

### IL-2v_5U_-T cells become polyfunctional TOX_neg_ effectors in tumors

We next asked how CD8^+^T cells adapt in the tumors following *in vitro* preconditioning with IL-2v versus IL-2. OT1-T cells expanded with IL-2v^5U^ were ∼80% PD1^neg^TCF-1^+^TOX^neg^ prior to ACT (**Figure 5H**). Strikingly, tumor-infiltrating IL-2v^5U^-OT1-T cells, in spite of upregulating PD-1 (i.e. indicating tumor recognition, consistent with tumor control) did not upregulate TOX after ACT, and transitioned from a PD1^+^TCF1^+^TOX^neg^ progenitor-like state (∼50% of cells on day 2 post-ACT) to a PD1^+^TCF1^neg^TOX^neg^ effector (TEFF)-like state (∼95% on day 14 post-ACT) (**Figure 5H** and **S6N**). Additionally, more than ∼90% of TCF1^neg^TOX^neg^ TEFF-like IL-2v^5U^-OT1-T cells were KLRG1^neg^ (**Figure S6O**), a marker for short-live effector cells found in acute infections and were functional as confirmed by restimulation ex *vivo* (**Figure 5I**). ScRNA-seq analysis of tumor-infiltrating IL-2v^5U^-OT1-T cells revealed (relative to baseline) important upregulation of TCR-activation markers *Cd137, Il2ra, Cd69* and *Ox40*, and TF_EFF_ regulon activity, consistent with intratumoral activation and upregulation of the effector program (**Figure 5J-5L**).

We have recently shown that CD8^+^T cells engineered to secrete constitutively IL-2v and IL-33, when transferred into non-lymphodepleted/irradiated tumor bearing mice, acquired a novel synthetic effector TOX^neg^ state (TSE) state transitioning through a synthetic precursor (TSP), and exhibited enhanced tumor control^46^. Projection of our scRNA-seq data from tumor-infiltrating IL-2v^5U^-OT1-T cells into a reference map including the canonical TOX^+^ progenitor-exhausted (TPEX) and terminally-exhausted (TEX) subsets and the novel TOX^neg^ synthetic states^46^ indicated marked overlap of the IL-2v-expanded TILs post-ACT with the synthetic states, with progression towards the TSE state by 10 days post-second ACT infusion (**Figure 5M** and **5N**). Thus, T cells expanded under IL-2v are stem-like cells with higher capacity for engraftment, expansion, and persistence in lymphodepleted hosts, which in tumors differentiate into TOX^neg^Klrg1^neg^ effector CD8^+^T cells without committing to the canonical exhaustion differentiation pathway, and they exhibit superior antitumor efficacy relative to T cells expanded under IL-2.

### IL-2v is a superior homeostatic cytokine for T-cell therapy

The use of IL-2 along with *γ*_c_ cytokines IL-15 and IL-7 has received great attention as an alternative to the IL-2 alone for ACT manufacturing, owing to greater frequencies of output T_CM_ cells, which persist more *in vivo* and achieve better tumor control^30, 47, 48^. Given the effects of IL-2v unveiled above, we examined whether expanding cells in IL-2v^5U^ plus IL-15 and IL-7 (hereafter referred to as IL-2v/15/7) enhances T-cell performance over IL-2v^5U^ alone. Interestingly, *in vitro* expansion with IL-2v/15/7 reduced by almost two-fold the overall OT1-T cell expansion relative to IL-2v, without altering the final proportion of TCM cells (**Figure 6A-6C**). Similarly, while the combination of IL-2 with IL-15+IL-7 enhanced memory differentiation compared to IL-2 alone, it also reduced total cell yield in culture compared to IL-2v. Additionally, the combination of IL-2v with IL-15+IL-7 did not improve *in vitro* effector cytokine/chemokine production or tumor cell killing, nor *in vivo* tumor control, mouse survival, nor persistence in tumor and spleen (**Figure 6D-6K**). Altogether, IL-2v is a superior homeostatic cytokine for expanding T cells for ACT, outperforming IL-2, IL-15, or IL-15+IL-7 for T-cell expansion due to its unique ability to promote the generation of large numbers of minimally differentiated CD8^+^ T cells with strong tumor control capability.

**Figure 6.**
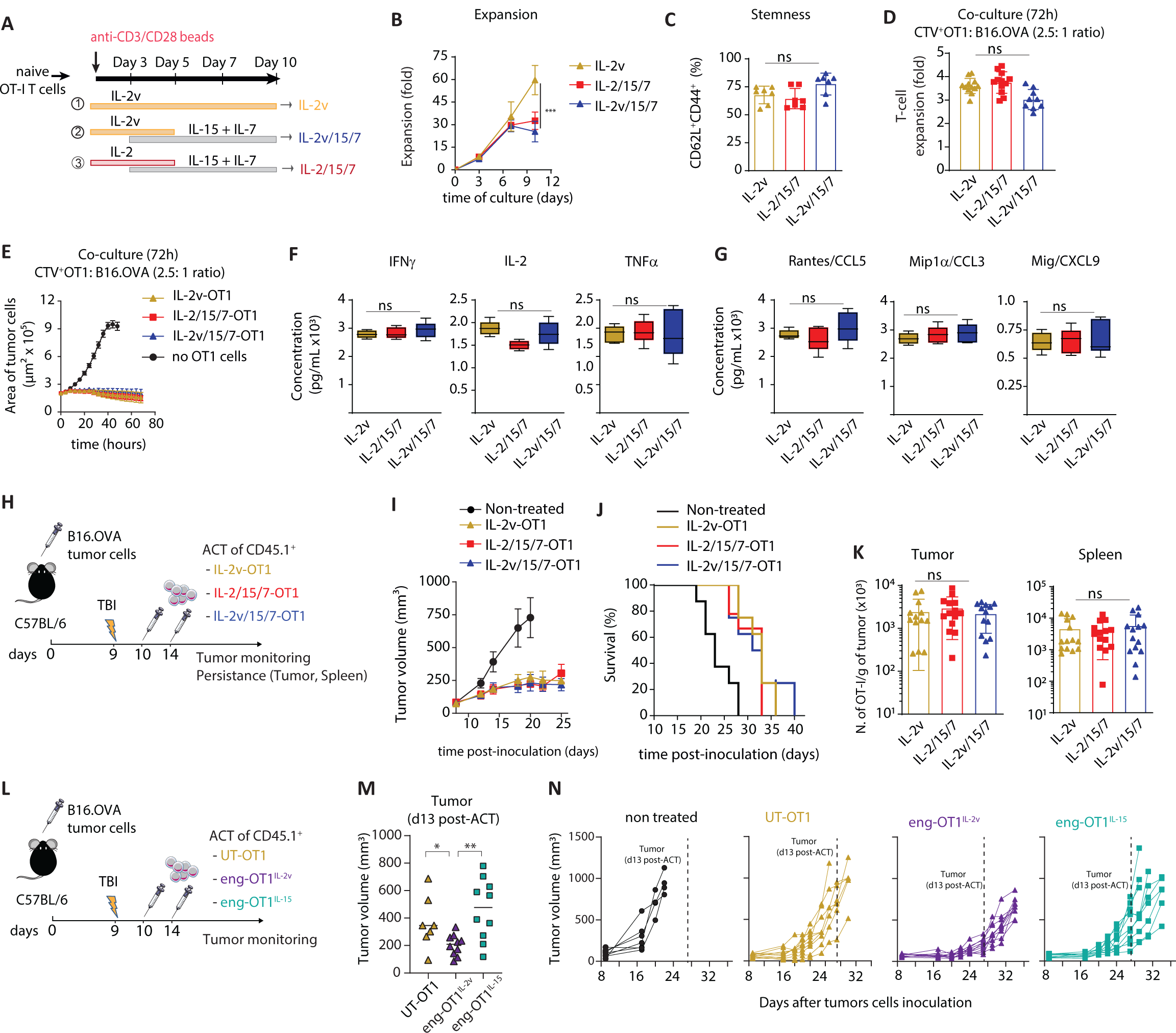
IL-2v is a superior homeostatic cytokine for T-cell therapy. (**A**) Schematic of naïve OT1-T cell activation and cytokine stimulation. (**B**) Kinetics of OT1-T cell expansion. (**C**) Frequency of CD62L^+^CD44^+^ OT1-T cells at day 10. (**D-G**) Expanded OT1-T cell (at day 10) co-culture with melanoma B16.OVA^NucLightRed^ tumor cells (in absence of exogenous cytokines). (**D**) OT1-T cell expansion at 72 hours of co-culture. (**E**) Growth kinetics of B16.OVA^NucLightRed^ tumor cells in co-culture, represented as the area of red fluorescent tumor cells detected in the IncuCyte live imaging platform. (**F**) Cytokine and (**G**) chemokine production after 24-hour co-culture. (**H**) Schematic of ACT study. (**I**) Tumor control and (**J**) survival curves. (**K**) Numbers of persisting OT1-T cells in the tumor and spleen at day 6 post-ACT. (**L**) Schematic of ACT *in vivo* study with T cells engineered to secrete IL-2v (eng-OT1^IL-2v^) versus IL-15 (eng-OT1^IL-15^). (**M**) Individual tumor control curves and (**N**) cumulative tumor volume on d13 post-ACT (corresponding to d27 after tumor cells inoculation). Data show mean ±SEM. For (**B-G** and **K**), n=7-12 biological replicates; pooled data from three independent experiments. For (**I**), n=8 biological replicates; representative of 3 independent experiments. For (**M** and **N**), n=7-10 biological replicates; representative data from 2 independent experiments. *p<0,05, **p<0,01 ***p<0,001, ****p<0,0001, and ns= non significative for p>0,05, using Unpaired t test with Welch’s correction.

Based on the above observations supporting the superiority of IL-2v for T-cell expansion for ACT, we sought to compare the efficacy of T cells engineered to secrete IL-2v (eng-OT1^IL-2v^) vs. IL-15 (eng-OT1^IL-15^) transferred in lymphodepleted (irradiated) B16.OVA-bearing mice. We observed better tumor control by eng-OT1^IL-2v^ and no significant impact of eng-OT1^IL-15^ relative to untransduced OT1-T cells (**Figure 6L-6N**). Thus, enforced IL-2v secretion by engineered CD8^+^T cells in the TME provides, in the context of preconditioning (sublethal irradiation), superior tumor control than IL-15 secretion.

## Discussion

The rational modification of cytokines to alter engagement with cognate receptors can be exploited to improve the biological properties of target cells. To date, however, the precise mechanisms by which this can be achieved have not been fully elucidated. Here, we have demonstrated that a non-IL-2R*α*-engaging IL-2v^26^ acquires IL-2R binding properties enabling the generation of optimal CD8^+^ T cells for ACT. While similar proliferation was observed for IL-2^50U^ and IL-2v^5U^ cultured T cells, an overall higher expansion was achieved for the variant due to increased viability of the cells. At the same time, a similar proportion of T_CM_ cells were generated as by IL-15. However, in contrast to the quiescent nature of T cells exposed to IL-15, we found that the memory/stem-like T cells generated by IL-2v^5U^ are ‘metabolically active’ and coined this hybrid state as T_MAS_. Thus, IL-2v coupled stemness to cell expansion and high metabolic activity, which further translated into superior effector potential *in vitro*, higher engraftment and expansion *in vivo*, and better tumor control by T cells cultured in IL-2v^5U^ vs. IL-2^50U^, as well as by T cells engineered with IL-2v vs. IL-15.

In order to elucidate the means by which IL-2v couples stemness to cell expansion and metabolic activity we took a systems-biology approach and dissected the signaling, gene regulatory networks and metabolic pathways at play. Overall, we found that the binding affinity of IL-2v /IL-2R and the dose of IL-2v used dictated the intensity and duration of IL-2v/IL-2R signaling, which in turn instructed specific TF programs. Specifically, we observed that low-intensity signals from IL-2v^5U^ or IL-15^50U^ rescued the downregulation of stemness-associated TF regulons initiated by TCR/CD28 activation of T cells. In contrast, the high-intensity signaling mediated by IL-2^50U^ drove the downregulation of stemness-associated TF regulons and promoted the activation of effector-associated TF regulons. Moreover, unlike IL-15^50U^, IL-2v^5U^ enabled sustained activation of proliferation-associated TF regulons, likely due to low but sustained/tonic signaling. Notably, functional annotation of the target genes of proliferation-associated TFs revealed involvement in cell cycle progression, chromosome segregation, and RNA and DNA metabolic processes. In line with this, for IL-2v^5U^ cultured T cells we observed an upregulation in fluxes of anabolic pathways including nucleotide production for RNA and DNA synthesis, the metabolism of structural cholesterol, glycerophospholipids and sphingolipids, and amino acid metabolism. In addition, IL-2v^5U^ expanded T cells exhibited increased capacity for mitochondrial ATP production without relying on aerobic glycolysis like IL-2^50U^ T cells.

Taken together, our work has important implications for development of IL-2 variants, which despite important preclinical promise have fallen short of expectations in the clinic^49, 50^. Here, we have demonstrated that a non-IL-2R*α*-engaging variant of IL-2 enables the generation and robust expansion of a novel metabolically active stem-like state of CD8^+^ T cells, which we coined T_MAS_ cells, having ideal properties for ACT. Importantly, we have also provided an experimental and systems-biology pipeline enabling the design and comprehensive evaluation of cytokines and their target cells with respect to function, signaling, gene-regulation and metabolic networks.

## Author Contributions

The study was supervised by G.C. and M.I.. G.C, M.I. and Y.O.M., conceived the project, designed the experiments, and interpreted results. Y.O.M., C.J.L., M.M.B., T.M., E.C., and J.C.O. performed experiments and interpreted results. Analysis of scRNA-seq datasets was performed by I.C., N.R. and M.M. Regulon analysis was performed by I.C. and N.R. K.L. contributed to supervision of early T-cell culture experiments performed by M.M.B. and E.C. and our observation that IL-2v favors a central memory phenotype. Fluxomic inference, analysis of Phospho-proteomics and metabolomics data was performed by M.M and supervised by V.H.. Seahorse and mitochondrial measurements were performed by H.C.H. and P.G., and supervised by N.V. Protein expression and purification was performed by B.S.. Western blots of phospho-proteins was performed by R.V.S, who also provided expertise on intracellular signaling.. Expert advice on T-cell metabolism was provided by P.C.H.. The overall manuscript was written and edited by G.C., M.I. and Y.O.M., with particular technical sections written by C.J.L., M.M., and I.C..

## Acknowledgments

We thank the R&D division of CIM (Havana, Cuba) for providing recombinant IL-2_V_. We thank Julien Marquis, Corinne Peter, and Karolina Bojkowska from Lausanne Genomic Technologies Facility at UNIL for the support with the scRNAseq experiments. We thank Francisco Sala de Oyanguren, Romain Bedel and Kevin Blackney from the Flow cytometry Facility at UNIL for the technical support. We thank Manfredo Quadroni and the Protein Analysis Facility at UNIL for the work on phospho-proteomics and Hector Gallart-Ayala and the UNIL Metabolomics Facility for their work.

## Funding

This work was generously supported by Ludwig Cancer Research, the Biltema and the ISREC Foundations, the Prostate Cancer Foundation, and an Advanced European Research Council Grant to G.C. (1400206AdG-322875), and the Swiss National Science Foundation to MI (SNSF# 310030_204326). C.J.L was partially supported by a postdoctoral fellowship from Ramon Areces Foundation.

## Data and materials availability

All data is available from the authors upon reasonable request.

## Data availability

scRNA-seq datasets analyzed during the current study are available in the GEO repository. Accession number: GSE223651 (https://www.ncbi.nlm.nih.gov/geo/query/acc.cgi?acc=GSE223651)

## Notes

https://www.ncbi.nlm.nih.gov/geo/query/acc.cgi?acc=GSE223651

### Competing Interest Statement

The authors have declared no competing interest.

## References

1. Boyman, O. & Sprent, J. The role of interleukin-2 during homeostasis and activation of the immune system. Nat Rev Immunol 12, 180–190 (2012).

2. Nelson, B.H., Lord, J.D. & Greenberg, P.D. Cytoplasmic domains of the interleukin-2 receptor beta and gamma chains mediate the signal for T-cell proliferation. Nature 369, 333–336 (1994).

3. Ponce, L.F., Montalvo, G., Leon, K. & Valiente, P.A. Differential Effects of IL2Ralpha and IL15Ralpha over the Stability of the Common Beta-Gamma Signaling Subunits of the IL2 and IL15 Receptors. J Chem Inf Model 61, 1913–1920 (2021).

4. Ross, S.H. & Cantrell, D.A. Signaling and Function of Interleukin-2 in T Lymphocytes. Annu Rev Immunol 36, 411–433 (2018).

5. Irving, M., Ortiz-Miranda, Y. & Coukos, G. IL-2 engineered MSCs rescue T cells in tumours. Nat Cell Biol 24, 1689–1691 (2022).

6. Lanitis, E., Dangaj, D., Irving, M. & Coukos, G. Mechanisms regulating T-cell infiltration and activity in solid tumors. Ann Oncol 28, xii18-xii32 (2017).

7. Gillis, S. & Smith, K.A. Long term culture of tumour-specific cytotoxic T cells. Nature 268, 154–156 (1977).

8. Morgan, D.A., Ruscetti, F.W. & Gallo, R. Selective in vitro growth of T lymphocytes from normal human bone marrows. Science 193, 1007–1008 (1976).

9. Rosenberg, S.A. IL-2: the first effective immunotherapy for human cancer. J Immunol 192, 5451–5458 (2014).

10. Kurnick, J.T., Kradin, R.L., Blumberg, R., Schneeberger, E.E. & Boyle, L.A. Functional characterization of T lymphocytes propagated from human lung carcinomas. Clin Immunol Immunopathol 38, 367–380 (1986).

11. Topalian, S.L., Muul, L.M., Solomon, D. & Rosenberg, S.A. Expansion of human tumor infiltrating lymphocytes for use in immunotherapy trials. J Immunol Methods 102, 127–141 (1987).

12. Topalian, S.L. et al. Immunotherapy of patients with advanced cancer using tumor-infiltrating lymphocytes and recombinant interleukin-2: a pilot study. Journal of clinical oncology: official journal of the American Society of Clinical Oncology 6, 839–853 (1988).

13. Rosenberg, S.A. & Restifo, N.P. Adoptive cell transfer as personalized immunotherapy for human cancer. Science 348, 62–68 (2015).

14. Rosenberg, S.A. & Dudley, M.E. Adoptive cell therapy for the treatment of patients with metastatic melanoma. Curr Opin Immunol 21, 233–240 (2009).

15. Mitchell, D.M., Ravkov, E.V. & Williams, M.A. Distinct roles for IL-2 and IL-15 in the differentiation and survival of CD8+ effector and memory T cells. J Immunol 184, 6719–6730 (2010).

16. Kalia, V. & Sarkar, S. Regulation of Effector and Memory CD8 T Cell Differentiation by IL-2-A Balancing Act. Front Immunol 9, 2987 (2018).

17. Kalia, V. et al. Prolonged interleukin-2Ralpha expression on virus-specific CD8+ T cells favors terminal-effector differentiation in vivo. Immunity 32, 91–103 (2010).

18. Pipkin, M.E. et al. Interleukin-2 and inflammation induce distinct transcriptional programs that promote the differentiation of effector cytolytic T cells. Immunity 32, 79–90 (2010).

19. Yao, X. et al. Levels of peripheral CD4(+)FoxP3(+) regulatory T cells are negatively associated with clinical response to adoptive immunotherapy of human cancer. Blood 119, 5688–5696 (2012).

20. Rohaan, M.W., van den Berg, J.H., Kvistborg, P. & Haanen, J. Adoptive transfer of tumor-infiltrating lymphocytes in melanoma: a viable treatment option. J Immunother Cancer 6, 102 (2018).

21. Mo, F. et al. An engineered IL-2 partial agonist promotes CD8(+) T cell stemness. Nature 597, 544–548 (2021).

22. Mitra, S. et al. Interleukin-2 activity can be fine tuned with engineered receptor signaling clamps. Immunity 42, 826–838 (2015).

23. Spangler, J.B. et al. Antibodies to Interleukin-2 Elicit Selective T Cell Subset Potentiation through Distinct Conformational Mechanisms. Immunity 42, 815–825 (2015).

24. Codarri Deak, L., et al. PD-1-cis IL-2R agonism yields better effectors from stem-like CD8(+) T cells. Nature 610, 161–172 (2022).

25. Quijano-Rubio, A. et al. A split, conditionally active mimetic of IL-2 reduces the toxicity of systemic cytokine therapy. Nat Biotechnol (2022).

26. Carmenate, T. et al. Human IL-2 mutein with higher antitumor efficacy than wild type IL-2. J Immunol 190, 6230–6238 (2013).

27. Carmenate, T. et al. The antitumor effect induced by an IL-2 ’no-alpha’ mutein depends on changes in the CD8(+) T lymphocyte/Treg cell balance. Front Immunol 13, 974188 (2022).

28. Condotta, S.A. & Richer, M.J. The immune battlefield: The impact of inflammatory cytokines on CD8+ T-cell immunity. PLoS Pathog 13, e1006618 (2017).

29. Wu, Z. et al. The IL-15 receptor alpha chain cytoplasmic domain is critical for normal IL-15Ralpha function but is not required for trans-presentation. Blood 112, 4411–4419 (2008).

30. Klebanoff, C.A. et al. IL-15 enhances the in vivo antitumor activity of tumor-reactive CD8+ T cells. Proc Natl Acad Sci U S A 101, 1969–1974 (2004).

31. Kaartinen, T. et al. Low interleukin-2 concentration favors generation of early memory T cells over effector phenotypes during chimeric antigen receptor T-cell expansion. Cytotherapy 19, 1130 (2017).

32. Gattinoni, L. et al. Acquisition of full effector function in vitro paradoxically impairs the in vivo antitumor efficacy of adoptively transferred CD8+ T cells. J Clin Invest 115, 1616–1626 (2005).

33. Masid, M., Ataman, M. & Hatzimanikatis, V. Analysis of human metabolism by reducing the complexity of the genome-scale models using redHUMAN. Nat Commun 11, 2821 (2020).

34. Pandey, V. & Hatzimanikatis, V. Investigating the deregulation of metabolic tasks via Minimum Network Enrichment Analysis (MiNEA) as applied to nonalcoholic fatty liver disease using mouse and human omics data. PLoS Comput Biol 15, e1006760 (2019).

35. Ring, A.M. et al. Mechanistic and structural insight into the functional dichotomy between IL-2 and IL-15. Nat Immunol 13, 1187–1195 (2012).

36. Levin, A.M. et al. Exploiting a natural conformational switch to engineer an interleukin-2 ‘superkine’. Nature 484, 529–533 (2012).

37. Martinez-Fabregas, J. et al. Kinetics of cytokine receptor trafficking determine signaling and functional selectivity. Elife 8 (2019).

38. Pais Ferreira, D., et al. Central memory CD8(+) T cells derive from stem-like Tcf7(hi) effector cells in the absence of cytotoxic differentiation. Immunity 53, 985–1000 e1011 (2020).

39. Tiemessen, M.M. et al. T Cell factor 1 represses CD8+ effector T cell formation and function. J Immunol 193, 5480–5487 (2014).

40. Kaech, S.M. & Cui, W. Transcriptional control of effector and memory CD8+ T cell differentiation. Nat Rev Immunol 12, 749–761 (2012).

41. Venet, D., Dumont, J.E. & Detours, V. Most random gene expression signatures are significantly associated with breast cancer outcome. PLoS computational biology 7, e1002240 (2011).

42. Nayar, R. et al. TCR signaling via Tec kinase ITK and interferon regulatory factor 4 (IRF4) regulates CD8+ T-cell differentiation. Proc Natl Acad Sci U S A 109, E2794–2802 (2012).

43. Iwata, A. et al. Quality of TCR signaling determined by differential affinities of enhancers for the composite BATF-IRF4 transcription factor complex. Nat Immunol 18, 563–572 (2017).

44. Rollings, C.M., Sinclair, L.V., Brady, H.J.M., Cantrell, D.A. & Ross, S.H. Interleukin-2 shapes the cytotoxic T cell proteome and immune environment-sensing programs. Sci Signal 11 (2018).

45. Hukelmann, J.L. et al. The cytotoxic T cell proteome and its shaping by the kinase mTOR. Nat Immunol 17, 104–112 (2016).

46. Corria-Osorio, J., Carmona, S. J., Stefanidis, E., Andreatta, M., Muller, T. Ortiz-Miranda, Y., Seijo, B., Castro, W., Jimenez-Luna, C., Scarpellino, L., Ronet, C., Spill, A., Lanitis, E., Luther, S.A., Romero, P., Irving, M., Coukos, G. Orthogonal Cytokine Engineering Enables Novel Synthetic Effector States in Tumor-rejecting CD8+ T Cells Escaping Canonical Exhaustion. Nature Immunology in press (2023).

47. Klebanoff, C.A. et al. Determinants of successful CD8+ T-cell adoptive immunotherapy for large established tumors in mice. Clin Cancer Res 17, 5343–5352 (2011).

48. Fraietta, J.A. et al. Biomarkers of Response to Anti-CD19 Chimeric Antigen Receptor (CAR) T-Cell Therapy in Patients with Chronic Lymphocytic Leukemia. Blood 128 (2016).

49. Mullard, A. Engineered IL-2 cytokine takes pivotal immuno-oncology blow. Nat Rev Drug Discov 21, 327 (2022).

50. Mullard, A. Restoring IL-2 to its cancer immunotherapy glory. Nat Rev Drug Discov 20, 163–165 (2021).

