## Supplemental Figures and captions for "Transcriptional reprogramming by IL-2 variant generates metabolically active stem-like T cells"

S.Figure 1

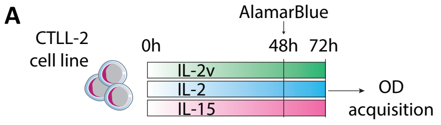


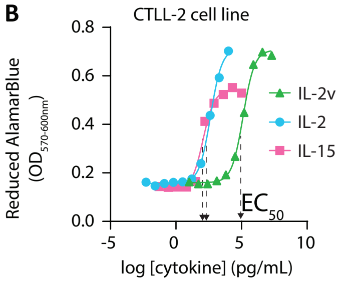


C

| cytokine | IU/mL | ng/mL |
|---|---|---|
| IL-2v | 1 | 130±20 |
| IL-2 | 1 | 0.2±0.1 |
| IL-15 | 1 | 0.2±0.1 |

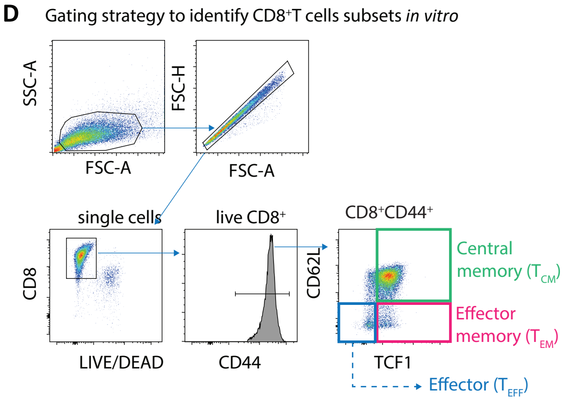

**Figure S1. IL-2v behaves as a weak IL-2R agonist and favors stemness and $CD8^+$T cell expansion *in vitro*. (A)** The CTLL-2 cell line was cultured with increasing doses of the indicated cytokine for 72 hours. Biological activity based on CTLL-2 cells proliferation was determined by assessment of AlamarBlue dye reduction. (**B**) Dose-response effect of IL-2v, IL-2 and IL-15 on CTLL-2 cells proliferation. (**C**) Concentration of the indicated cytokine equivalent to 1IU/mL of activity determined by $EC_{50}$ values obtained in (**B**). (**D**) Gating strategy for the identification of $T_{CM}$, $T_{EM}$ and $T_{EFF}$ phenotypes among $CD8^+CD44^+$ T cells. Data show mean ±SEM. For **(B** and **C**), n= 3 technical replicates; representative data from 3 independent experiments. $T_{CM}$, central memory; $T_{EM}$, effector memory; $T_{EFF}$, effector $CD8^+$T cells.

S.Figure 2

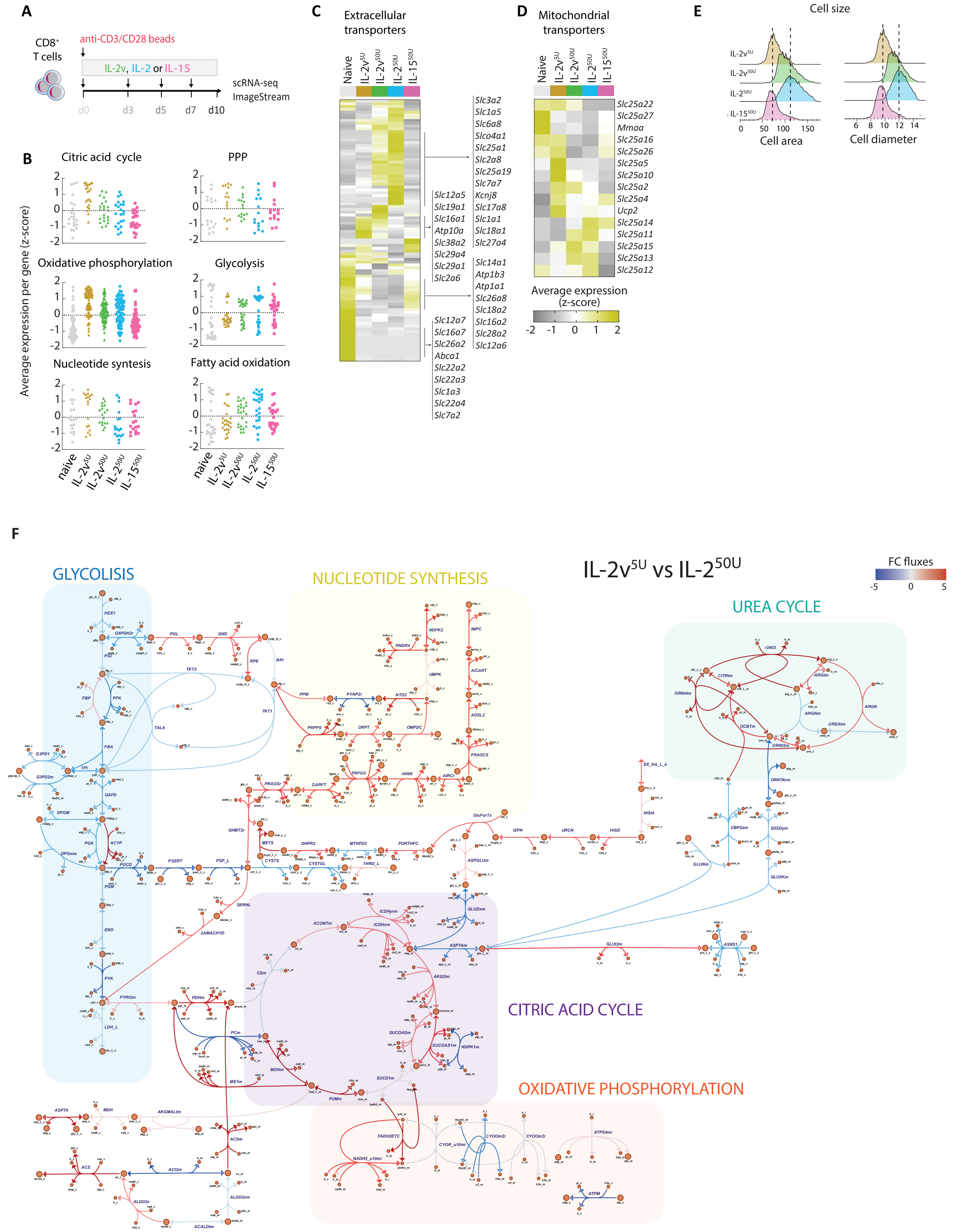
A
CD8+ T cells
anti-CD3/CD28 beads
IL-2v, IL-2 or IL-15
d0
d3
d5
d7
d10
scRNA-seq
ImageStream
B
Average expression per gene (z-score)
Citric acid cycle
PPP
Oxidative phosphorylation
Glycolysis
Nucleotide syntesis
Fatty acid oxidation
naive
IL-2v5U
IL-2v50U
IL-250U
IL-1550U
C
Extracellular transporters
Naive
IL-2v5U
IL-2v50U
IL-250U
IL-1550U
Slc3a2
Slc1a5
Slc6a8
Slco4a1
Slc25a1
Slc2a8
Slc25a19
Slc7a7
Slc12a5
Slc19a1
Slc16a1
Atp10a
Slc38a2
Slc29a4
Slc29a1
Slc2a6
Kcnj8
Slc17a8
Slc1a1
Slc18a1
Slc27a4
Slc14a1
Atp1b3
Atp1a1
Slc26a8
Slc18a2
Slc16a2
Slc28a2
Slc12a6
Slc12a7
Slc16a7
Slc26a2
Abca1
Slc22a2
Slc22a3
Slc1a3
Slc22a4
Slc7a2
D
Mitochondrial transporters
Slc25a22
Slc25a27
Mmaa
Slc25a16
Slc25a26
Slc25a5
Slc25a10
Slc25a2
Slc25a4
Ucp2
Slc25a14
Slc25a11
Slc25a15
Slc25a13
Slc25a12
Average expression (z-score)
E
Cell size
IL-2v5U
IL-2v50U
IL-250U
IL-1550U
Cell area
Cell diameter
F
GLYCOLISIS
NUCLEOTIDE SYNTHESIS
IL-2v5U vs IL-250U
FC fluxes
UREA CYCLE
CITRIC ACID CYCLE
OXIDATIVE PHOSPHORYLATION

**Figure S2. IL-2v alters CD8$^+$T cells metabolic gene expression/fluxes and cell size compared to IL-2.** (**A**) Schematic of naïve OT1-T cell activation and cytokine stimulation during expansion for scRNA-seq analysis on day 10. (**B**) OT1-T cell expression of genes corresponding to the indicated metabolic pathways. (**C** and **D**) Heatmaps showing average gene expression of (**C**) extracellular and (**D**) mitochondrial transporters at day 10. (**E**) Cell size evaluated by imaging flow cytometry (ImageStream). scRNA-seq, single cell RNA sequencing; PPP, pentose phosphate pathway. (**F**) Fold change in average metabolic fluxes in IL-2v$^{5U}$-OT1-T cells versus IL-2$^{50U}$ -OT1-T cells after 10 days of expansion (See Methods for details).

S.Figure 3

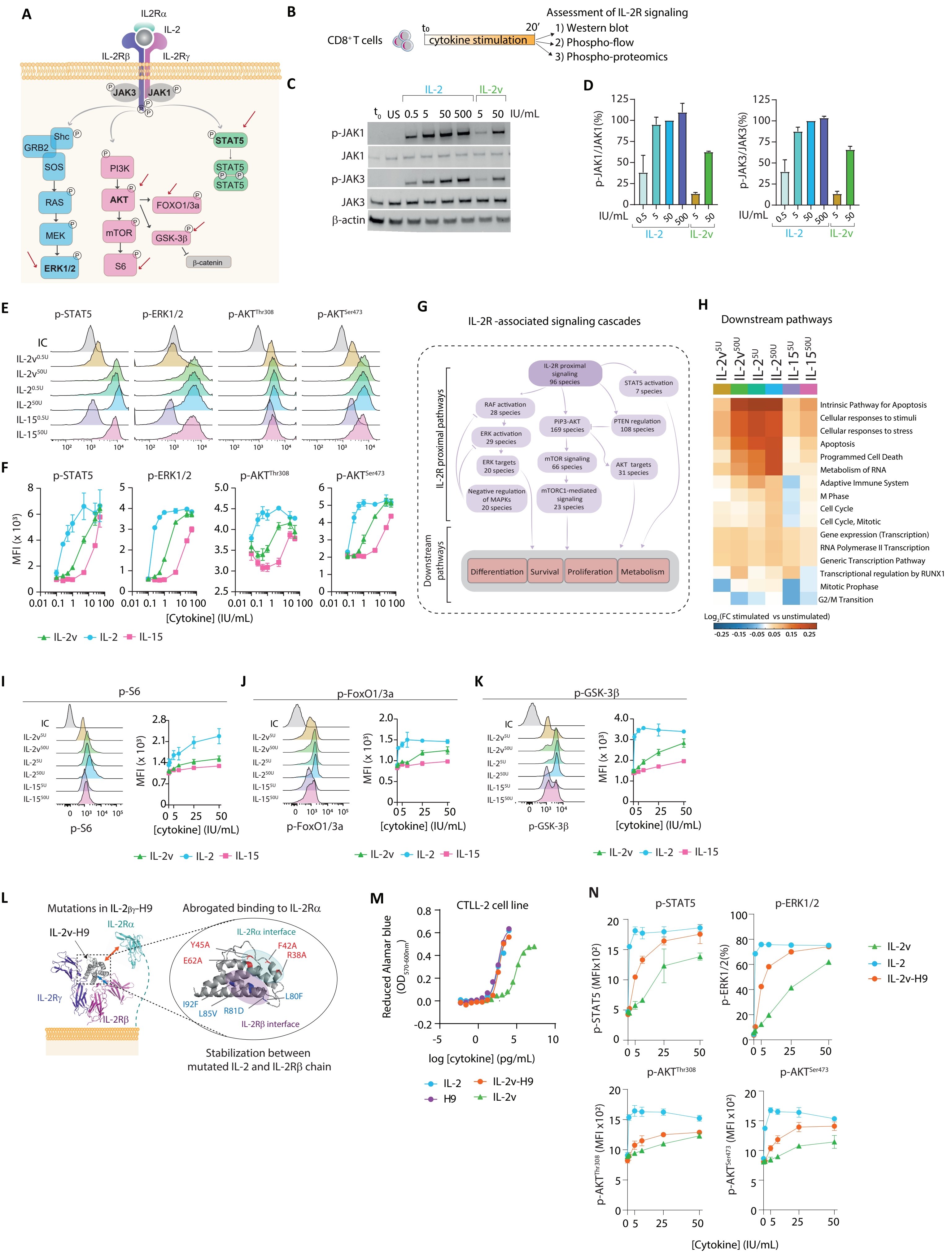

A
IL2Rα
IL-2
IL-2Rβ
IL-2Rγ
JAK3
JAK1
Shc
GRB2
SOS
RAS
MEK
ERK1/2
PI3K
AKT
mTOR
S6
FOXO1/3a
GSK-3β
β-catenin
STAT5
B
CD8+ T cells
t0
20′
cytokine stimulation
Assessment of IL-2R signaling
1) Western blot
2) Phospho-flow
3) Phospho-proteomics
C
IL-2
IL-2v
t0 US 0.5 5 50 500 5 50 IU/mL
p-JAK1
JAK1
p-JAK3
JAK3
β-actin
D
p-JAK1/JAK1(%)
p-JAK3/JAK3(%)
IU/mL
IL-2
IL-2v
E
p-STAT5
p-ERK1/2
p-AKT Thr308
p-AKT Ser473
IC
IL-2v 0.5U
IL-2v 50U
IL-2 0.5U
IL-2 50U
IL-15 0.5U
IL-15 50U
F
MFI (x 10³)
[Cytokine] (IU/mL)
IL-2v
IL-2
IL-15
G
IL-2R -associated signaling cascades
IL-2R proximal pathways
Downstream pathways
IL-2R proximal signaling 96 species
STAT5 activation 7 species
RAF activation 28 species
ERK activation 29 species
ERK targets 20 species
Negative regulation of MAPKs 20 species
PIP3-AKT 169 species
PTEN regulation 108 species
mTOR signaling 66 species
AKT targets 31 species
mTORC1-mediated signaling 23 species
Differentiation
Survival
Proliferation
Metabolism
H
Downstream pathways
IL-2v 5U
IL-2v 50U
IL-2 5U
IL-2 50U
IL-15 5U
IL-15 50U
Intrinsic Pathway for Apoptosis
Cellular responses to stimuli
Cellular responses to stress
Apoptosis
Programmed Cell Death
Metabolism of RNA
Adaptive Immune System
M Phase
Cell Cycle
Cell Cycle, Mitotic
Gene expression (Transcription)
RNA Polymerase II Transcription
Generic Transcription Pathway
Transcriptional regulation by RUNX1
Mitotic Prophase
G2/M Transition
Log2(FC stimulated vs unstimulated)
-0.25 -0.15 -0.05 0.05 0.15 0.25
I
p-S6
MFI (x 10³)
[cytokine] (IU/mL)
J
p-FoxO1/3a
K
p-GSK-3β
L
Mutations in IL-2βγ-H9
Abrogated binding to IL-2Rα
IL-2v-H9
IL-2Rα
IL-2Rγ
IL-2Rβ
IL-2Rα interface
Y45A
F42A
R38A
E62A
I92F
L85V
R81D
L80F
IL-2Rβ interface
Stabilization between mutated IL-2 and IL-2Rβ chain
M
CTLL-2 cell line
Reduced Alamar blue (OD570-600nm)
log [cytokine] (pg/mL)
IL-2
H9
IL-2v-H9
IL-2v
N
p-STAT5
p-ERK1/2
p-AKT Thr308
p-AKT Ser473
p-STAT5 (MFIx10²)
p-ERK1/2(%)
p-AKT Thr308 (MFI x10²)
p-AKT Ser473 (MFI x10²)
[Cytokine] (IU/mL)
IL-2v
IL-2
IL-2v-H9

**Figure S3. IL-2v induces in mouse and human CD8$^+$T cells lower-intensity IL-2R signaling than IL-2 that is critical for memory commitment.** (**A**) Schematic of proximal IL-2R signaling pathways. (**B**) CD8$^+$T cell stimulation with the indicated cytokine prior to western blot (**c** and **d**), phospho-flow cytometry (**E, F,** and **I-K**) or phospho-proteomics (**H**). (**C**) Representative western blot of JAK1/3 phospho-proteins in stimulated OT1-T cells, and (**D**) mean phosphorylation status of JAK1/3 proteins relative to its total level normalized to the condition IL-2$^{50U}$. (**E**) Representative histograms and (**F**) dose-response effect of IL-2v, IL-2 or IL-15 on the phosphorylation levels of the indicated signaling molecules in human CD8$^+$T cells. (**G**) Schematic of IL-2R signaling network including the number of phospho-species per sub-pathway used for phospho-proteomic analysis (refer to Methods for details). (**H**) Activity of IL-2R distal pathways based on phospho-site enrichment analysis of stimulated OT1-T cells. (**I-K**) Representative histograms (left) and dose-response effect (right) of IL-2v, IL-2 or IL-15 on the phosphorylation levels of the indicated signaling molecules. **(L)** Mutations introduced in IL-2v-H9 variant. (**M**) CTLL-2 cell line was cultured with increasing doses of IL-2v, IL-2v-H9 or IL-2 for 72 hours. Proliferation (biological activity) was determined based on AlamarBlue dye reduction. (**N**) Dose-response effect of IL-2v, IL-2v-H9 or IL-2 on the phosphorylation levels of the indicated species in OT1-T cells. Data show mean ±SEM. For **(D**), n=3 biological replicates; representative data from 2 independent experiments. For (**F**), n= 6-9 donors; representative data from 3 independent experiments. For (**I-K** and **N**), n=3 biological replicates; representative of 2-3 independent experiments. For (**H**), n= 3 biological replicates. *p<0,05, **p<0,01 ***p<0,001, ****p<0,0001 using Unpaired t test with Welch's correction. For (**M**), n= 3 technical replicates; representative data from 3 independent experiments.

S.Figure 4

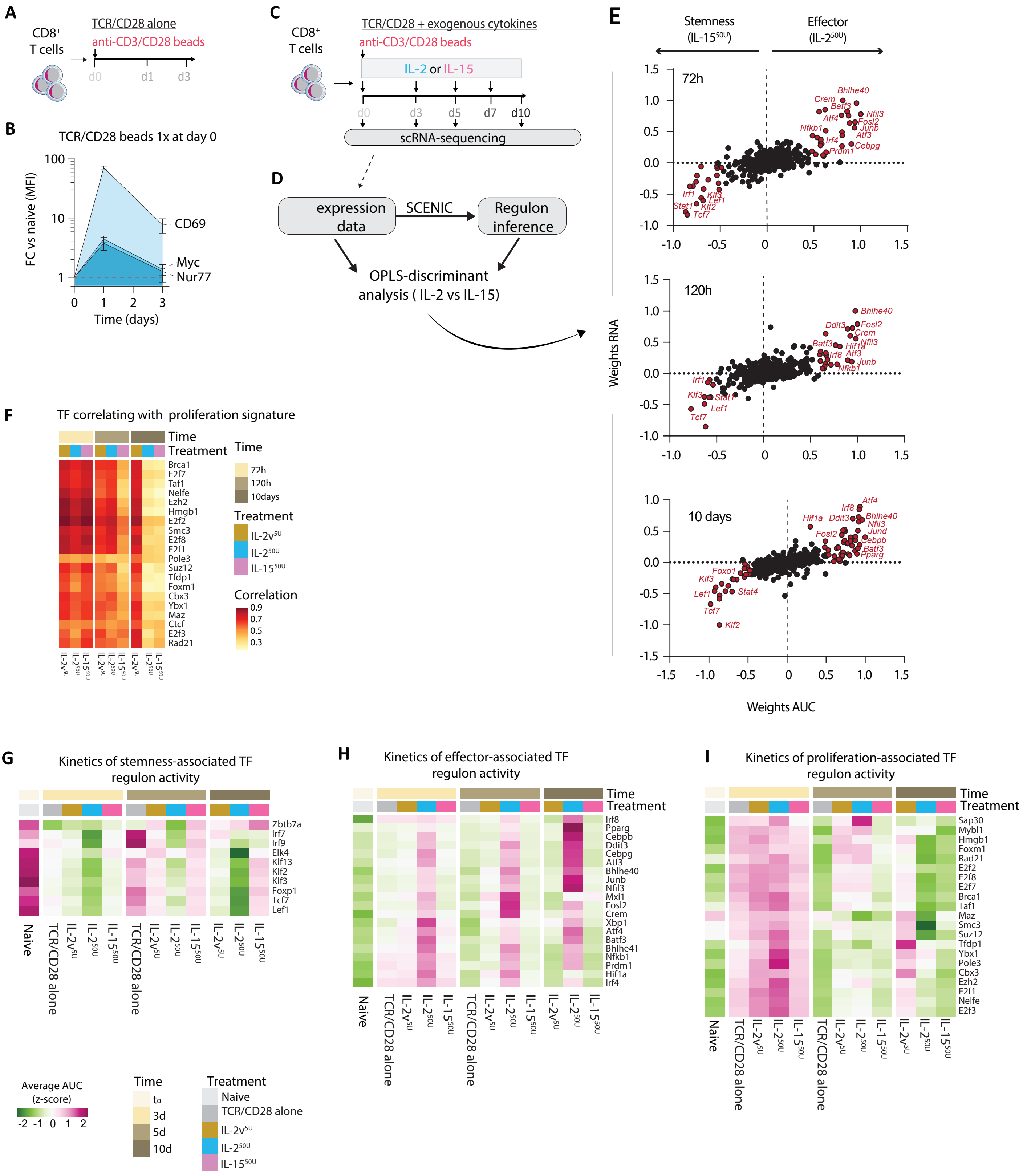
A
CD8+ T cells
TCR/CD28 alone
anti-CD3/CD28 beads
d0
d1
d3
B
TCR/CD28 beads 1x at day 0
FC vs naive (MFI)
CD69
Myc
Nur77
Time (days)
C
CD8+ T cells
TCR/CD28 + exogenous cytokines
anti-CD3/CD28 beads
IL-2 or IL-15
d0
d3
d5
d7
d10
scRNA-sequencing
D
expression data
SCENIC
Regulon inference
OPLS-discriminant analysis ( IL-2 vs IL-15)
E
Stemness (IL-15 50U)
Effector (IL-2 50U)
72h
120h
10 days
Weights RNA
Weights AUC
F
TF correlating with proliferation signature
Time
Treatment
72h
120h
10days
Correlation
G
Kinetics of stemness-associated TF regulon activity
H
Kinetics of effector-associated TF regulon activity
I
Kinetics of proliferation-associated TF regulon activity
Naive
TCR/CD28 alone
IL-2v 5U
IL-2 50U
IL-15 50U
Average AUC (z-score)
Time
t0
3d
5d
10d
Treatment

**Figure S4. Effects of IL-2v$^{5U}$, IL-2$^{50U}$ or IL-15$^{50U}$ on the kinetics of the activation of regulons linked to CD8$^{+}$T cell differentiation and proliferation.** (**A**) Schematic of naïve OT1-T cells activation and expansion. (**B**) Kinetics of CD69, Myc and Nur77 expression levels relative to naïve OT1-T cells as evaluated by flow cytometry. n=9, pooled data from 3 independent experiments. (**C**) Schematic of naïve OT1-T cell activation and cytokine stimulation for up to 10 days. Samples were collected at the indicated time points to perform scRNA-seq. (**D**) Schematics of OPLS discriminant analysis. Both TFs expression and regulon activity from IL-2$^{50U}$ or IL-15$^{50U}$-OT1-T cells were used to perform the analysis (refer to Methods for details). (**E**) Ranking of TFs based on their discriminant power for IL-2$^{50U}$ or IL-15$^{50U}$-OT1-T cells at the indicated time points. TFs in red represent those with the highest ranking. (**F**) Heatmap showing TFs with the highest correlation scores to cell proliferation genes (from metaPCNA signature, Table S1) across treatments and time points (refer to Methods for details). OPLS, Orthogonal Projections to Latent Structures. (**G-I**) Heatmaps showing average regulon activity (AUC) for (**G**) stemness-, (**H**) effector- and (**I**) proliferation-associated regulons in OT1-T cells stimulated with TCR/CD28 stimulation alone or plus the indicated cytokine for up to 10 days.

S.Figure 5

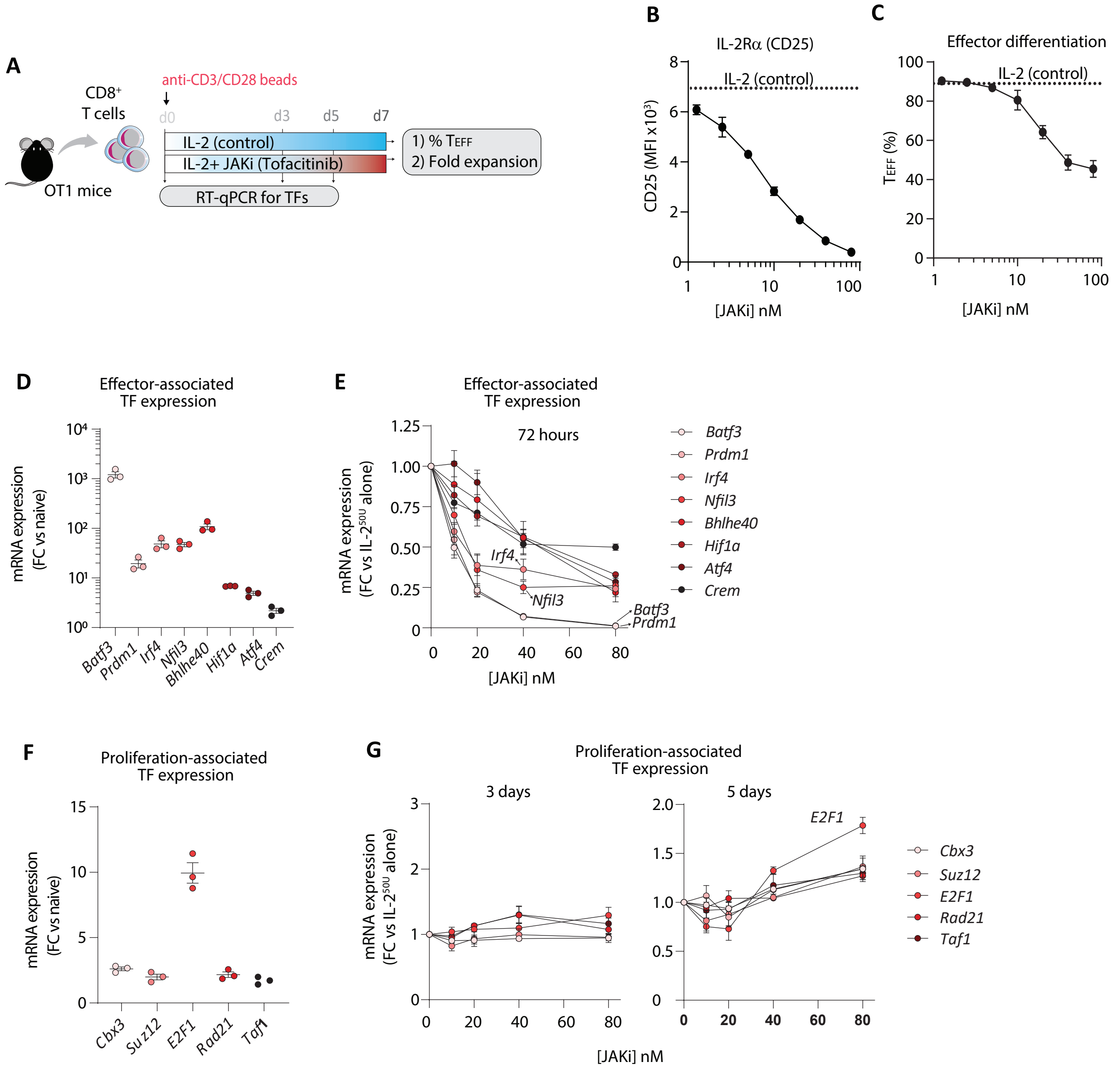

A
anti-CD3/CD28 beads
CD8+ T cells
d0 d3 d5 d7
IL-2 (control)
IL-2+ JAKi (Tofacitinib)
1) % TEFF
2) Fold expansion
OT1 mice
RT-qPCR for TFs
B
IL-2Rα (CD25)
IL-2 (control)
CD25 (MFI x10³)
[JAKi] nM
C
Effector differentiation
IL-2 (control)
TEFF (%)
[JAKi] nM
D
Effector-associated TF expression
mRNA expression (FC vs naive)
Batf3 Prdm1 Irf4 Nfil3 Bhlhe40 Hif1a Atf4 Crem
E
Effector-associated TF expression
72 hours
mRNA expression (FC vs IL-2 50U alone)
Irf4
Nfil3
Batf3
Prdm1
[JAKi] nM
Batf3 Prdm1 Irf4 Nfil3 Bhlhe40 Hif1a Atf4 Crem
F
Proliferation-associated TF expression
mRNA expression (FC vs naive)
Cbx3 Suz12 E2F1 Rad21 Taf1
G
Proliferation-associated TF expression
3 days
5 days
E2F1
mRNA expression (FC vs IL-2 50U alone)
[JAKi] nM
Cbx3 Suz12 E2F1 Rad21 Taf1

**Figure S5. Effects of attenuated IL-2R signal strength on effector- and proliferation-associated TFs expression.** (**A**) Schematic of naïve OT1-T cells activation and expansion in the presence of IL-$2^{50U}$ +/- various doses of JAK inhibitor (JAKi=Tofacitinib) for 7 days. (**B** and **C**) Dose-response effect of JAK inhibition on (**B**) CD25 expression and (**C**) frequency of $T_{EFF}$ cells within OT1-T cells at day 7. (**D** and **F**) Fold change of (**D**) effector- and (**F**) proliferation-associated TFs expression in OT1-T at day 3 versus naïve OT1-T cells. (**E** and **G**) Dose-response effect of JAK inhibition on (**E**) effector- and (**G**) proliferation-associated TFs expression in OT1-T at the indicated time points. Expression values were normalized to the condition IL-$2^{50U}$. Data show mean ±SEM. n=3 biological replicates; representative data from 2 independent experiments. AUC, activity unit per cell; $T_{EFF}$, effector $CD8^{+}$T cells; and FC, fold change.

### S. Figure 6

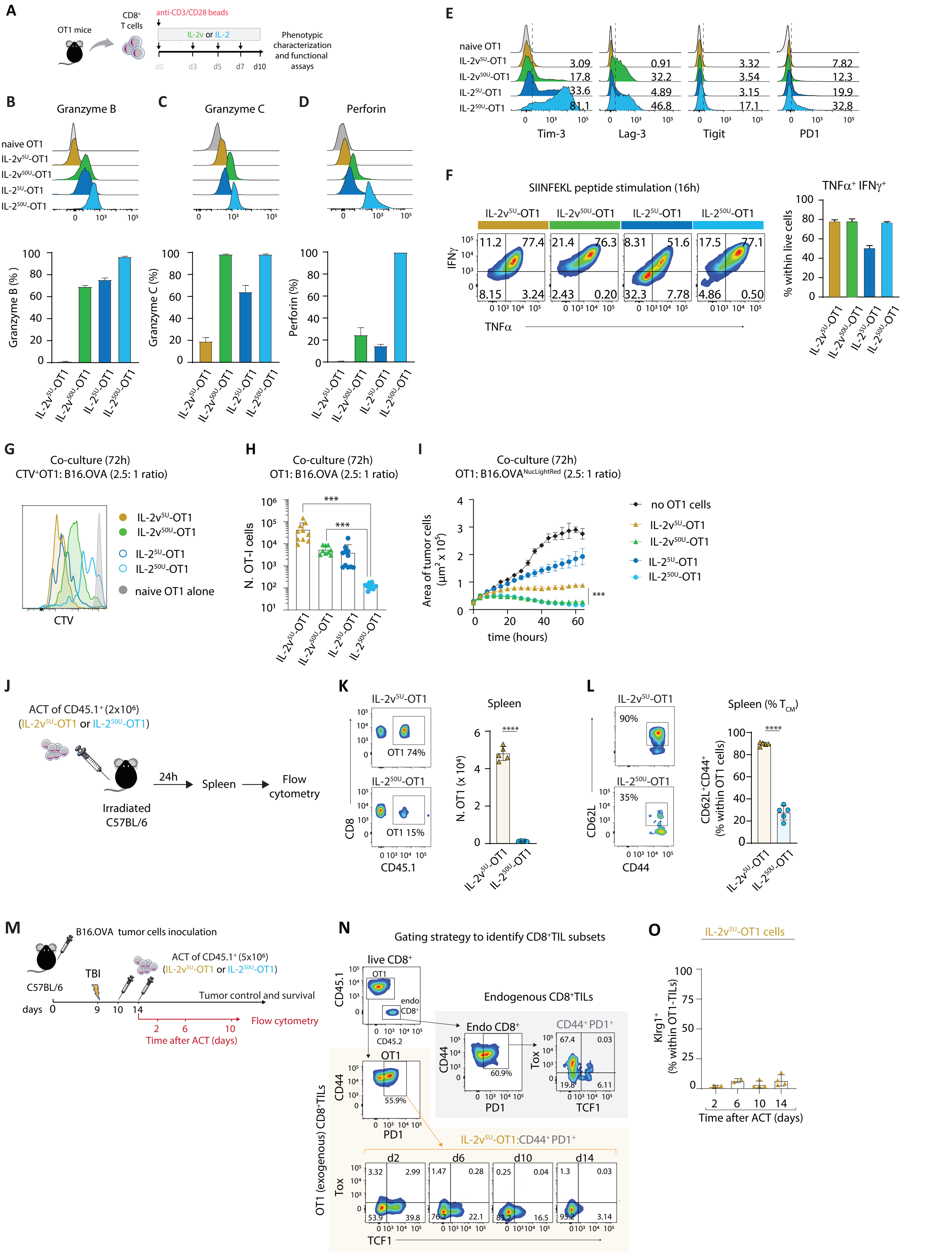

A
OT1 mice
CD8+ T cells
anti-CD3/CD28 beads
IL-2v or IL-2
d0 d3 d5 d7 d10
Phenotypic characterization and functional assays
B
Granzyme B
C
Granzyme C
D
Perforin
naive OT1
IL-2v5U-OT1
IL-2v50U-OT1
IL-25U-OT1
IL-250U-OT1
Granzyme B (%)
Granzyme C (%)
Perforin (%)
E
Tim-3
Lag-3
Tigit
PD1
F
SIINFEKL peptide stimulation (16h)
TNFα+ IFNγ+
IFNγ
TNFα
% within live cells
G
Co-culture (72h)
CTV+OT1: B16.OVA (2.5: 1 ratio)
naive OT1 alone
CTV
H
Co-culture (72h)
OT1: B16.OVA (2.5: 1 ratio)
N. OT-I cells
I
Co-culture (72h)
OT1: B16.OVANucLightRed (2.5: 1 ratio)
no OT1 cells
Area of tumor cells (μm2 x 105)
time (hours)
J
ACT of CD45.1+ (2x106)
(IL-2v5U-OT1 or IL-250U-OT1)
24h
Spleen
Flow cytometry
Irradiated C57BL/6
K
Spleen
OT1 74%
OT1 15%
CD8
CD45.1
N. OT1 (x 104)
L
Spleen (% TCM)
CD62L
CD44
CD62L+CD44+ (% within OT1 cells)
M
B16.OVA tumor cells inoculation
C57BL/6
TBI
ACT of CD45.1+ (5x106)
(IL-2v5U-OT1 or IL-250U-OT1)
Tumor control and survival
days 0 9 10 14
Flow cytometry
Time after ACT (days)
N
Gating strategy to identify CD8+TIL subsets
live CD8+
CD45.1
CD45.2
endo CD8+
Endogenous CD8+TILs
Endo CD8+
CD44+ PD1+
Tox
TCF1
PD1
CD44
OT1 (exogenous) CD8+TILs
IL-2v5U-OT1:CD44+ PD1+
d2 d6 d10 d14
O
IL-2v5U-OT1 cells
Klrg1+ (% within OT1-TILs)
Time after ACT (days)

**Figure S6. Functional profile of CD8$^{+}$T cells expanded in IL-2v compared to IL-2.** (**A**) Schematic of naïve OT1-T cell activation and cytokine stimulation during expansion for functional analyses on day 10. Representative histograms (**top**) and cumulative frequency (**bottom**) of (**B**) Granzyme B, **(C)** Granzyme C, and **(D)** perforin expression by OT1-T cells. (**E**) Representative histograms of inhibitory receptor expression by OT1-T cells. (**F**) Representative dot plots (**left**) and cumulative frequency (**right**) of co-production of IFNγ and TNFα by OT1-T cells upon re-stimulation with SIINFEKL peptide. (**F-H**) Co-culture of OT1-T cells with melanoma B16.OVA$^{NucLightRed}$. (**G**) OT1-T cell proliferation as evaluated CTV-dilution and (**H**) overall OT1-T cell expansion at 72 hours of co-culture. (**I**) Growth kinetics of B16.OVA$^{NucLightRed}$ tumor cells in co-culture, represented as the area of red fluorescent tumor cells detected in the IncuCyte live imaging platform. (**J**) Schematic of ACT study. (**K**) Representative dot plots (**left**) and cumulative data (**right**) of the number of OT1-T-cells in the spleen 24-hours after transfer as indicated in (**J**). (**L**) Representative dot plots (**left**) and cumulative frequencies (**right**) of $T_{CM}$ cells among OT1-T cells in the spleen 24-hours after transfer as indicated in (**J**). (**M**) Schematic of ACT study. Tumor and/or TDLN tissues were collected at the indicated time points after second ACT infusion for flow cytometric analyses. (**N**) Representative dot plots of the phenotype of IL-2v$^{5U}$-OT1 TILs post-ACT relative to endogenous CD8$^{+}$TILs. (**O**) Proportion of Klrg1$^{+}$ IL-2v$^{5U}$-OT1 cells post ACT. Data show mean ±SEM. For (**B-F** and **I**), n=4 biological replicates, representative of 2-3 independent experiments. For (**H**), n=8-10 biological replicates; pooled from 3 independent experiments. For (**K, L** and **O**), n=3-5 biological replicates, representative of 2-3 independent experiments. *p<0,05, **p<0,01 ***p<0,001, ****p<0,0001 using Unpaired t test with Welch's correction. CTV, CellTrace Violet; ACT, adoptive T cell therapy; TILs, tumor infiltrating lymphocytes; $T_{CM}$, central memory CD8$^{+}$T cells.
