## Supplementary material for "Transcriptional reprogramming by IL-2 variant generates metabolically active stem-like T cells": Materials and Methods

### Reagents or resources

| REAGENT or RESOURCE | SOURCE | IDENTIFIER |
|---|---|---|
| **Antibodies** | | |
| FCR blocking reagent (clone 2.4G2) | BD Biosciences | Cat# 553141 |
| PE/Cy5.5 anti-mouse CD3e (clone 2. 145-2C11) | Invitrogen | Cat# 14-0031-82 |
| Brilliant Violet 711 anti-mouse CD8a (clone 2. 53-6.7) | Biolegend | Cat# 100748 |
| PE/Dazlee 594 anti-mouse CD45.1 (clone A20) | Biolegend | Cat# 110748 |
| PE-mouse anti-human IgG4 (clone HP6025) | abcam | Cat# 99825 |
| Brilliant Violet - anti-mouse/human CD44 (clone IM7) | Biolegend | Cat# 103040 |
| APC/Cy7-anti-mouse PD-1 (clone 29F.1A12) | Biolegend | Cat# 135224 |
| APC- anti-mouse Granzyme C (clone SFC1D8) | Biolegend | Cat# 150812 |
| PE/Cy7- anti-mouse Granzyme C (clone SFC1D8) | Biolegend | Cat# 150804 |
| Rabbit anti TCF1 (TCF7) antibody (clone C63D9) | Cell Signaling Technology | Cat# 2203S |
| PE-anti-rabbit IgG (H+L), F(ab')2 Fragment | Cell Signaling Technology | Cat # 8885S |
| Alexa Fluor 488-anti-rabbit IgG (H+L), F(ab')2 Fragment | Cell Signaling Technology | Cat # 4408S |
| PE/Cy7- anti-mouse Granzyme B (clone CLB-GB11) | Novus Biological | Cat# NBP1-50071PECY7 |
| Brilliant Violet 421 anti-mouse TIM-3(clone RMT3-23) | Biolegend | Cat# 119723 |
| PE/Cyanine7 anti-mouse TIGIT Antibody (clone Vstm3)) | Biolegend | Cat# 142107 |
| APC anti-mouse CD223 (LAG-3) Antibody (C9B7W) | Biolegend | Cat# 125210 |
| Brilliant Violet 605 anti-mouse/human KLRG1(clone 2F1/KLRG1) | Biolegend | Cat# 138419 |
| PerCP-Cyanine5.5- anti-mouse IFNg (clone XMG1.2) | Invitrogen | Cat# 45-7311-82 |
| PE/Cy7- anti-mouse TNFa (clone MP6-XT22) | Biolegend | Cat# 506324 |
| PE/Cy7- anti-mouse CD62L (clone MEL-14) | Biolegend | Cat# 104418 |
| PE- anti-TOX antibodies human and mouse (clone REA473) | Miltenyi | Cat# 130-107-785 |
| APC- Armenian Hamster IgG Isotype Ctrl Antibody (clone HTK888) | Biolegend | Cat# 400912 |
| PE/Cy7- Armenian Hamster IgG Isotype Ctrl Antibody (clone HTK888) | Biolegend | Cat# 400922 |
| APC-Perforin (S16OO9A) | Biolegend | Cat# 154304 |
| PercPCy5.5-CD45.2 | Biolegend | Cat# 109828 |
| Alexa Fluor 488 Donkey anti-Rat IgG (H+L) Highly Cross-Adsorbed Secondary Antibody | Invitrogen | Cat#A21-208 |
| InVivoMAb anti-mouse CD25 (IL-2Rα) (clone PC-61.5.3) | Bio X Cell | Cat# BE0012 |

| | | |
|---|---|---|
| Phospho-p44/42 MAPK (Erk1/2) (Thr202/Tyr204) (197G2) Rabbit mAb | Cell Signaling Technologies | Cat# 4377 |
| Phospho-Akt (Thr308) (244F9) Rabbit mAb | Cell Signaling Technologies | Cat# 4056 |
| Phospho-Akt (Ser473) (D9E) XP Rabbit mAb | Cell Signaling Technologies | Cat# 4060 |
| Phospho-FoxO1 (Thr24)/FoxO3a (Thr32)/FoxO4 (Thr28) (4G6) Rabbit mAb | Cell Signaling Technologies | Cat# 2599 |
| Phospho-GSK-3β (Ser9) (D85E12) XP® Rabbit mAb | Cell Signaling Technologies | Cat# 5558 |
| Phospho-Stat5 (Tyr694) (C11C5) Rabbit mAb | Cell Signaling Technologies | Cat# 9359 |
| Phospho-Jak1(Tyr1034/1035) (D7N4Z) Rabbit mAb | Cell Signaling Technologies | Cat# 74129 |
| Jak1 (6G4) Rabbit mAb | Cell Signaling Technologies | Cat# 3344 |
| Phospho-Jak3 (Tyr980/981) (D44E3) Rabbit mAb | Cell Signaling Technologies | Cat# 5031 |
| Jak3 (D7B12) Rabbit mAb | Cell Signaling Technologies | Cat# 8863 |
| β actin | Santa Cruz | Cat# sc-47778 |
| **Bacterial and Virus Strains** | | |
| Replication-defective MSGV retrovirus | produced in house | |
| Competent E.coli | produced in house | |
| **Biological Samples** | | |
| **Chemicals, Peptides, and Recombinant Proteins** | | |
| Live/Dead Fixable Aqua Dead cell stain Kit | ThermoFisher Scientific | Cat# L34957 |
| Recombinant human IL-2 Variant | Produced in house | |
| Recombinant human IL-2 | Peprotech | Cat# 200-02-1mg |
| Recombinant human IL-15 | Miltenyi | Cat# 130-095-766 |
| Recombinant human IL-7 | Miltenyi | Cat# 130-095-361 |
| Brefeldin A | Enzo Life Sciences | Cat# BML-G405-0005 |
| Turbofect | Thermo Fisher Scientific | Cat# 12172220 |
| MitoTracker™ Green FM | (Thermo Fisher Scientific) | Cat# M7514 |
| Glucose | Sigma-Aldrich | Cat# G8270 |
| pyruvate | Thermo Fisher Scientific | Cat# 11360070 |

| | | |
|---|---|---|
| glutamine | Thermo Fisher Scientific | Cat# 25030024 |
| oligomycin | Sigma-Aldrich | Cat# O4876-5MG |
| carbonyl cyanide-p-trifluoromethoxyphenylhradone (FCCP) | Sigma-Aldrich | Cat# C2920-10mg |
| rotenone | Sigma-Aldrich | Cat# R8875-1g |
| antimycin A | Sigma-Aldrich | Cat# A8674-25mg |
| CellTrace Violet | Thermo Fisher Scientific | Cat# C34557 |
| alamarBlue™ HS Cell Viability Reagent | Thermo Fisher Scientific | Cat# A50100 |
| Recombinant Retronectin | Takara | Cat# T202 |
| Tofacitinib, Janus kinase (JAK) inhibitor | Abcam | Cat# ab142068 |
| CellEvent Caspase-3/7 Green Detection Reagent | Thermo Fisher Scientific | Cat# R37111 |
| **Critical Commercial Assays** | | |
| Tumor Dissociation Kit | Miltenyi | Cat# 130-096-730 |
| JETSTAR 2.0 Plasmid Maxiprep Kit | Genomed | Cat# 210050 |
| Fixation/Permeabilization solution kit | BD Biosciences | Cat# 554714 |
| Foxp3/Transcription factor staining Buffer Set | ThermoFisher Scientific | Cat# 00-5523-00 |
| EasySep™ Mouse T Cell Isolation Kit | Stemcell Technologies | Cat# 19851 |
| EasySep™ Human CD8+ T Cell Isolation Kit | Stemcell Technologies | Cat# 17953 |
| mouse CD45 MicroBeads | Miltenyi | Cat# 130-052-301 |
| Dynabeads Mouse T-Activator CD3/CD28 | Gibco | Cat# 11452D |
| **Deposited Data** | | |
| RNAseq data | generated in this Study | To be deposited soon in GEO |
| **Experimental Models: Cell Lines** | | |
| Mouse: B16-OVA (derived from B16-F10:RRID:CVCL_0159) | Generated in house | |
| Mouse: B16-OVA NucLightRed | Generated in house | |
| Phoenix™- Eco retrovirus producer lines | Invitrogen | Cat# 18324012 |
| **Experimental Models: Organisms/Strains** | | |
| C57BL/6 mice | Charles River Laboratory | Strain 027 |
| Mouse: B6.Tg (TcraTcrb)1100Mjb/J (OT-1transgenic CD45.1) | (Hogquist et al.1994) P.Romero (UNIL) | RRID :IMSR_JAX :03831 |

### Cell lines and culture conditions

Ovalbumin (OVA)-expressing B16-F10 murine melanoma (B16.OVA) cell lines were cultured in Dulbecco's Modified Eagle's Media (DMEM) supplemented with 10% fetal bovine serum (FBS), 100 U/mL of penicillin, and 100 μg/mL streptomycin sulphate (Gibco, Thermo Fisher Scientific). B16.OVA cell line stably expressing nuclear-localized mKate2 (B16.OVA$^{NucLight Red}$), a red fluorescent protein, were generated by transduction using commercial lentivirus (Lenti Nuclight-Red, Incucyte, Sartorius) according to the manufacturer's instructions. The murine cytotoxic T lymphocyte CTLL-2 and Phoenix-eco retroviral ecotropic packaging cell lines were maintained in RPMI 1640-Glutamax media (Gibco) supplemented with 10% FBS, 100 U/mL penicillin and 100 μg/mL streptomycin sulfate. Cell lines were routinely tested for mycoplasma contamination. Primary human and murine T cells were cultured in RPMI 1640-Glutamax media supplemented with 10% FBS, 100 U/mL penicillin, 100 μg/mL streptomycin sulfate and cytokines as described in the experiments. Also, murine T cells media included 1mM Pyruvate and 50 μM β-mercaptoethanol.

### Mouse strains

Mice were housed at the University of Lausanne (UNIL, Epalinges, Switzerland) animal facility. All *in vivo* experiments were conducted in accordance and approval from the Service of Consumer and Veterinary Affairs (SCAV) of the Canton of Vaud. Female C57BL/6 mice aged 6-8 weeks were purchased from Harlan (Harlan, Netherlands) and CD45.1$^+$ OT1 TCR mice (C57BL/6-Tg(TcraTcrb)1100Mjb/J) were bred in-house.

### Production and purification of IL-2v and other IL-2 variants

IL-2v (E62A, Y45A, F42A, R38A) was kindly provided by the Development Facility of the Center for Molecular Immunology (CIM; Habana, Cuba). The H9 superkine (I92F, L85V,

R81D, and L80F), additional IL-2v, and the IL-2v-H9 double mutein were produced in-house. Genes were synthesized by GeneArt (Thermo Fisher Scientific) and cloned in a Pet22b bacterial expression vector. Rosetta pRAREII competent cells were transformed with the expression vectors, plated and single colonies isolated (Amp-resistant) and amplified in LBAC medium (LB Ampicillin 150mg/ml-Chloramphenicol 34mg/mL). Bacterial pellets were washed and lysed by 3-step sonication. Aggregates were then denatured under SDS-based conditions, followed by addition of $CuSO_4$ solution to promote disulfide bridge formation and KCl for precipitation of the SDS. Finally, the IL-2 variants were purified by size exclusion chromatography.

**CTLL-2 proliferation assay to evaluate the biological activity of IL-2 and variants**

The biological activity of IL-2, IL-2v, H9 superkine and IL-2v-H9 was determined using a dye reduction proliferation assay of CTLL-2 cells as previously described (1). Briefly, CTLL-2 cells were washed three times and cultured at $2x10^5$ cells/mL in the presence of various doses of IL-2 or the variants for 48 hours. AlamarBlue dye (Thermo Fisher Scientific) was then added, and the cells cultured for a further 16-20 hours. Absorbance at 570-600 nm was measured to evaluate cell proliferation. $EC_{50}$ (half maximal effective concentration) of the dose-response curves was used to determine the biological activity of the variants.

**Primary immune cell isolation**

Spleens from $CD45.1^+$ OT1 mice were mechanically homogenized by filtering through a 70 µm cell strainer. Red blood cells were lysed with ammonium chloride potassium buffer. Human peripheral blood mononuclear cells (PBMCs) from healthy donors (buffycoats preparation) were obtained by a standard protocol of centrifugation based on Lymphoprep (Axonlab) separation solution. Murine and human $CD8^+$ T cells were purified by negative magnetic selection using the EasySep™ Mouse T Cell Isolation Kit and EasySep™ Human $CD8^+$ T Cell Isolation Kit (Stemcell Technologies), respectively, according to the manufacturer's protocol. For *ex vivo* evaluation of immune cell infiltrate, lymph nodes and B16.OVA tumors, at described time-points post-ACT, were mechanically dissociated, and passed through a 70-µm cell strainer.

#### *In vitro* T cell activation and expansion

Purified $CD8^+$ T cells were plated at $0.5x10^6$ cells/mL density in 24- or 48-well plates in T-cell medium supplemented with wild-type recombinant human IL-2 (IL-2; Peprotech), IL-2v or recombinant human IL-15 (IL-15; Miltenyi) at the indicated doses. Mouse and human T cells were activated on day 0 with anti-CD3/CD28 antibody coated Dynabeads (Gibco) at a ratio of 2 beads. Alternatively, $CD8^+$ T cells were cultured in presence of IL-2 or IL-2v until day 5, combined with IL-15 and IL-7 (Miltenyi) at 10 ng/mL each from day 3 onwards. Finally, at day 10 and/or 14 cells were used for subsequent analysis. In indicated experiments, $CD8^+$ OT1 cells were expanded with IL-2 plus 10 µg/mL of anti-mouse CD25 mAb (PC-61, BioXcell) or IL-2v-H9 at the indicated doses. For the experiments involving JAK inhibition, $CD8^+$OT1 cells were activated as above and cultured in presence of IL-2 (50 IU/mL) with or without the JAK inhibitor, Tofacitinib (Abcam), at different doses up to 7 days. Fold T cell expansion was calculated by dividing the absolute number of live T cells by the number of cells plated at the beginning of the culture (day 0).

#### Retroviral constructs and virus preparation

Retrovirus was produced and concentrated as described in Lanitis et al. (2). Briefly, Phoenix Eco cells were seeded at $1 \times 10^7$ per T-150 tissue culture flask in 25mL culture medium 24-hours prior to transfection with 14.4 ug of pCL-Eco Retrovirus Packaging Vector and 21.4 ug of pMSGV transfer plasmid (coding for IL-2v or human IL-15) using Turbofect (Thermo Fisher Scientific). All plasmids were purified using JETSTAR 2.0 Plasmid Maxiprep Kit (Genomed). For the transfection mixture, a 3:1 ratio of turbofect:plasmid was prepared in 2ml of Optimem and incubated for 30 minutes at RT. Medium was then removed from T-150 flasks bearing 80-90% confluent Phoenix Eco cells and the transfection mixture was applied and incubated for 1 minute, followed by addition of 25 mL fresh medium. The viral supernatant was harvested 48 hours post-transfection followed by addition of 25 ml of fresh media. A second harvest was done again 24 hours later. The viral particles in both SN were concentrated by ultracentrifugation for 2-hours at 24,000g at 40C with a Beckman JS-24 rotor (Beckman Coulter) and suspended in 0.5 mL murine T-cell medium, then viral titer was determined using the method described by Lanitis et al. Finally, the retrovirus was aliquoted, frozen on dry ice and stored at -80°C.

**T cell transduction**

Untreated 24-well plates were coated with 20 µg/mL of recombinant retronectin (Takara) at 4°C for 24 hours and pre-loaded with 250µL retroviral supernatants by centrifugation at 2,000g for 90 min at 32ºC. Subsequently, 0.5. x $10^6$ of 24 hours activated $CD8^+$OT1 T cells were plated per well and incubated overnight. The next day, the transduction was repeated, and the T cells were cultured for an additional 5 days. The cultures were maintained at a cell density of 0.5-1x$10^6$ cells/mL and replenished with fresh T-cell media every 2-3 days supplemented with 50 IU/mL of IL-2.

**Antibodies and flow cytometric analysis**

For phenotypic analysis of $CD8^+$OT1 T cells and TILs the following fluorochrome-conjugated antibodies were used: Granzyme B (GB11), Granzyme C (SFC1D8) LAG-3 (C9B7W), TIGIT

(1G9), PD1 (29F.1A12), Perforin (S16009A), CD25 (PC-61), TNFα (MP6XT22), TIM-3 (RMT3-23), CD45.1 (A20), CD45 (30F/11), CD62L (MEL-14), CD44 (IM7), CD3ε (145-2C11), and CD8α (53-6.7), KLRG1 (2F1/KLRG1), from BioLegend; IFNγ (XMG1.2) from eBioscience; TOX (REA473) from Miltenyi Biotec, TCF-1 (C63D9) and polyclonal anti-rabbit from Cell Signaling Technology. LIVE/DEAD™ fixable aqua death cell staining (Thermo Fisher Scientific) was performed according to the manufacturer's instructions to exclude dead cells. Fc receptor binding was blocked by pre-incubating cells with purified rat anti-mouse CD16/CD132 (mouse BD Fc block). Cell-surface staining with antibodies was performed for 30 min at 4˚C. Intracellular cytokines, granzyme B, and TCF-1 staining were performed using the Foxp3/Transcription Factor Staining Buffer Set (Thermo Fisher Scientific) according to manufacturer's recommendations. For intracellular cytokine detection, T cells were also previously incubated with Brefeldin A (10μg/mL, Enzo Life Sciences) in standard culture conditions for 4 hours. For apoptosis evaluation, caspase activity was determined using CellEvent Caspase-3/7 Green Detection Reagent (R37111; Thermo Fisher Scientific) according to manufacturer's instructions. For mitochondria analysis, $CD8^+$OT1 cells were stained 10 days post *in vitro* culture by MitoTracker green (Thermo Fisher Scientific) and tetramethylrhodamine methyl ester (TMRM; Thermo Fisher Scientific), according to manufacturer's instructions. Cells were acquired with an LSRII flow cytometer (BD Bioscences). Additionally, we determined the cell diameter and area of OT1 T cells expanded for 10 days in presence of IL-2V2v (5 and 50 IU/mL), IL-2 (50 IU/mL) or IL-15 (50 IU/mL) by using an Amnis ImageStreamX Mk (Amnis, Merck Millipore). Data were analyzed using FlowJo software (TreeStar, Inc).

**Seahorse analysis**

For analysis of the metabolic phenotype of $CD8^+$OT1 T cells, oxygen consumption rate (OCR) and extracellular acidification rate (ECAR) were measured using a Seahorse XF96 Extracellular Flux Analyzer. T cells were plated in a Cell-Tak coated Seahorse microplate ($10^5$ cells/well) and preincubated at 37 ˚C for 45 min in the absence of $CO_2$. Seahorse media was supplemented with 10 mM glucose (Sigma-Aldrich), 1mM pyruvate (Thermo Fisher Sicentific) and 2 mM glutamine (Thermo Fisher Sicentific). ECAR was measured under basal conditions while OCR was measured under basal conditions and after the addition of the following drugs: oligomycin (1 μM), carbonyl cyanide-p-trifluoromethoxyphenylhradone (FCCP; 1.5 μM) and rotenone (0.5 μM) and antimycin A (0.5 μM; all from Sigma-Aldrich), following standard mitostress protocol.

**Proliferation assay**

Naïve $CD8^+$OT1 cells were stained with 5 μM CellTrace™ Violet (CTV, Thermo Fisher Scientific) according to manufacturer's recommendations. The cells were then activated with anti-CD3/anti-CD28 coated beads and expanded *in vitro* for 7 days in the presence of IL-2, IL-2V or IL-15 (5, 10, 25 and 50 IU/mL). Alternatively, in indicated experiments, $CD8^+$T cells were cultured in the presence of IL-2v and IL-2 (5 and 50 IU/mL) for 10 days and stained with 5 μM CTV (Thermo Fisher Scientific). The frequency of proliferating T cells was assessed as the percentage of cells that experienced at least one cellular division by flow cytometry based on CTV dilution.

**Cytokine detection assays**

$CD8^+$ OT1 T cells were co-cultured with B16.OVA cells at 2.5:1 (effector: target) ratio for 24 hours, supernatants collected and released cytokines and chemokines quantified using

Mouse Cytometric Bead Array (CBA) kit (BD Biosciences) as per the manufacturer's protocol. Expanded OT1 T cells (day 10), were stimulated with plate bound anti-CD3 (1μg/mL) for 4 hours in the presence of Brefeldin A (10μg/mL, Enzo Life Sciences). T cells were subsequently harvested, stained with antibodies to quantify intracellular IFNγ and TNFα levels by flow cytometry.

**Cytotoxicity assay**

*In vitro* tumor killing assays were performed using the IncuCyte ZOOM imaging platform (Essen Bioscience). Tumor cells were plated in 96-well plates one day before co-culture with T cells. Briefly, B16.OVA$^{NucLight\ Red}$ cells were co-cultured with $CD8^+$ OT1 T cells previously expanded *in vitro* for 10 days, at a 2.5:1 (effector: target) ratio. Tumor cells alone were used as a negative control of cell death. Images were captured every 2 hours for 72 hours and red living target cells were quantified with the IncuCyte ZOOM integrated analysis software.

**scRNA-Seq analysis**

**Sample preparation**

Naïve $CD8^+$OT1 T cells were activated *in vitro* on day 0 with anti-CD3/anti-CD28 coated beads and expanded in presence of IL-2, IL-2v or IL-15 (5 and/or 50 IU/mL) up to 10 days. Next, cells at different time points of *in vitro* culture (before expansion, day 0 and after 3, 5, 7, and 10 days) were processed for scRNAseq analysis. Sample processing and sequencing was performed by the Lausanne Genomic Technologies Facility, Center for Integrative Genomics, University of Lausanne. In addition, $CD8^+$OT1 TILs previously cultured with IL-2v (5 IU/mL) either untransduced, engineered to produced IL-2v or IL-15, and then used for ACT, were studied by scRNAseq 2 or 10 days after the therapy. For scRNA-seq of $CD8^+$OT1

TILs, live $CD44^+CD45.1^+CD8^+T$ cells were sorted using a BD FACSAria™ III (BD Bioscences).

**Data processing of scRNA-seq Libraries**

Cell Ranger count (10X Genomics, version 3.0.2) was used for the alignment and quantification of the scRNA-seq reads against the GRCh38 reference genome. All additional analyses were performed using R 4.0.2 and Seurat v3.0 1, unless otherwise indicated.

**Single-Cell Data Filtering and Normalization**

To filter out low-quality transcriptomes, total ribosomal and mitochondrial count filters were applied, with maximum accepted percentage counts coming from ribosomal and mitochondrial genes of 60% and 10%, respectively. In addition, the acceptable number of detected genes per cell ranges between 250 and 6000.

After the filtering, data was normalized using the “LogNormalize” function from the “Seurat v3.0.2” package, which normalizes the feature expression measurements for each cell by the total expression, multiplies this by a scale factor of 10,000 and log-transforms the result.

**Single-Cell Data dimensionality reduction and visualization**

Scaled z-scores for each gene were calculated using the ScaleData function from the “Seurat v3.0.2” and regressed against the number of UMIs per cell. The resulting scaled data was used as the input of a principal component analysis (PCA) on the top 1,000 most variable genes. The first 30 principal components were used to generate uniform manifold approximation and projection (UMAP) projections, with a minimum distance of 30 neighbors, unless it is indicated otherwise.

### Regulon analysis

#### Regulon inference

Regulons were inferred using the SCENIC pipeline (https://scenic.aertslab.org) 2. To detect transcriptional regulatory events that only take place in some experimental conditions, we split the dataset and carried out the analysis for each treatment. SCENIC integrates three algorithms (grnBoost2, RcisTarget and AUCell) corresponding to three sequential steps, and we used them as follows:

Step 1: Identifying co-expression modules using grnBoost2, which is a faster implementation of the original Genie3 algorithm 3, and SCENIC R package. The aggregation of targets into raw putative regulons was done using the runSCENIC_1_coexNetwork2modules function from the SCENIC R package with nTopTfs and nTopTargets parameters set to 50 and 5, respectively.

Step 2: Regulons and GRN inference. Co-expression modules (raw putative regulons, i.e., sets of genes regulated by the same transcription factor) are refined by removing indirect targets by motif discovery analysis using cisTarget algorithm and a cis-regulatory motif database 4,5. At this step, we used mm9-500bp-upstream-7species.mc9nr.feather and mm9-tss-centered-10kb-7species.mc9nr.feather cisTarget databases, and the motifs-v9-nr.mgi-m0.001-o0.0.tbl motif database. The motif database includes a score for each pair motif-gene, which allows the generation of a motif-gene ranking. A motif enrichment score is then calculated for the list of transcription factor selected targets by calculating the Area Under the recovery Curve (AUC) on the motif-gene ranking 2 using the RcisTarget R package (https://github.com/aertslab/RcisTarget). If a motif is enriched among the list of transcription factor targets, a regulon is derived including the target genes with a high motif-gene score.

Step 3: Finally, AUCell was used to quantify the regulon activity in each individual cell (https://github.com/aertslab/AUCell). AUCell provides a AUC score for each regulon and cell; we discarded regulons with less than 5 constituent elements, as the estimation of the activity of small regulons is less reliable.

**OPLS-DA**

In this work, we used the ropls R package6 to perform the Orthogonal Partial Least Squares discriminant analysis (OPLS-DA). The OPLS-DA is a multivariate analysis that exploits the pre-known data structure (classes) to remove the intra- class dispersion and maximize the inter- class dispersion, to derive a predictive axis. The resulting predictive axis constitute a projection of the regulon activity matrix that optimizes the split between the elements (cell) of the two given pre-known classes (experimental conditions, i.e., treatments). The absolute value of the relative contribution of each feature (regulon) to the predictive axis (weightStarMN value) was normalized between 0-1 preserving the sign of the original value. The resulting number become the discriminant score of the regulon. Within the context of a given OPLS-DA, the resulting discriminant score for the features included in the analysis can be compared, and a feature ranking can be generated. The input data for the OPLS-DA was scaled (mean centered and divided by the standard deviation).

**Connectivity analysis**

For the connectivity analysis we used the gene regulatory network (GRN) derived from the assembly of all regulons partial networks inferred in the regulon analysis. In order to test whether the number of regulatory interactions within a given sent of genes were significantly higher than expected by chance from a random selection of TFs, a connectivity analysis is performed using the SANTA algorithm 7. The SANTA algorithm calculates the area under the Knet function curve (AUK), which is computed for the observed gene set and for permutations of the same number of TFs. An empirical p-value is calculated using a Z-test. In this work, we used 10,000 permutations for each test and considered as significant p-values <= 0.05.

**Regulon clustering**

Transcription factors were clustered by regulon activity using the hierarchical clustering function embedded within the pheatmap R package (https://rdrr.io/cran/pheatmap/).

The unsupervised hierarchical clustering was performed using Euclidean distances and complete (complete linkage) as clustering method. The cutree_rows argument, which defines the number of clusters was set to 12. As the input for the clustering, we aggregated the regulon activity matrix taking, for each transcription factor, the mean regulon activity value for each of the experimental conditions (treatments) and scaled (mean centered and divided by the standard deviation) the resulting matrix.

**RT-qPCR**

Total RNA from T cells was isolated using RNeasy Mini Kit (Qiagen). For quantitative real-time PCR (RT-qPCR) studies, 1 ng of total RNA was reverse-transcribed using PrimeScript First Strand cDNA Synthesis Kit (Takara), as per manufacturer's instructions. Expression levels of target genes were determined using SYBR Green Fast PCR Master Mix (Applied Biosystems). Primer sequences are provided in Table 1. Expression was normalized to Ubiquitin C expression. All assays were performed using a 7500 Fast Real-Time PCR System (Applied Biosystems). Fold changes in expression were calculated by the ΔΔCt method.

*Table 1. Primers used for RT-qPCR*

| Probe Name | Proble code |
|---|---|
| Taqman primers mouse_ Cbx3 | Mm00850539_g1 |
| Taqman primers mouse_ Taf1 | Mm01229174_m1 |
| Taqman primers mouse_ Suz12 | Mm01304152_m1 |
| Taqman primers mouse_ E2f1 | Mm00432939_m1 |
| Taqman primers mouse _Rad21 | Mm00485474_m1 |
| Taqman primers mouse_Hif1a | Mm00468869_m1 |
| Taqman primers mouse_Prdm1 | Mm00476128_m1 |
| Taqman primers mouse_Nfil3 | Mm00600292_s1 |
| Taqman primers mouse_Batf3 | Mm01318274_m1 |
| Taqman primers mouse_Atf4 | Mm00515325_g1 |
| Taqman primers mouse_Bhlhe40 | Mm00478593_m1 |
| Taqman primers mouse_Irf4 | Mm00516431_m1 |
| Taqman primers mouse_UBC | Mm02525934_g1 |

### Metabolic fluxomic inference

### Reconstruction of a metabolic model for T-cells

We generated a reduced model around the metabolic subsystems that are of interest for the study of T cells. To this end, we applied the redHUMAN method (3) to the human genome-scale metabolic network Recon 3D (4). We used the composition of the RPMI medium to define the uptakes in the model, and we allowed also all the inorganic metabolites to be uptaken or secreted in the model. We selected 15 starting subsystems, namely, glycolysis/gluconeogenesis, citric acid cycle, pentose phosphate pathway, arginine and proline metabolism, pyruvate metabolism, glutamate metabolism, lycine, serine, alanine, and threonine metabolism, methionine and cysteine metabolism, urea cycle, ROS detoxification, purine synthesis, pyrimidine synthesis, glutathione metabolism, oxidative phosphorylation, and all the mitochondrial reactions. We used the redHUMAN parameters, Smin for redGEMX, D=1 for redGEM and Sminp3 for lumpGEM. As a result, we reconstructed redTCELL, a metabolic model with 967 metabolites, 1080 genes and 2151 reactions associated to 63 metabolic subsystems (Annexed Table 1).

**Transcriptomics data integration to generate context-specific CD8$^+$ T cell models.**

Based on the experimental data, we assumed a maximum doubling time of 6h for T cells treated with IL-2$^{50U}$, IL-2v$^{50U}$, and IL-2v$^{5U}$, and a maximum doubling time of 8h for T cells treated with IL-15$^{50U}$.

Moreover, we identified in the scRNAseq data a total of 1387 metabolic genes present in the Recon 3D model. For each T cell treatment (IL-2$^{50U}$, IL-2v$^{50U}$, IL-2v$^{5U}$, and IL-15$^{50U}$), we generated pseudo-bulk RNAseq data by averaging the corresponding scRNA-seq data for the metabolic genes, and we evaluated the gene-protein reaction (GPR) rules present in the metabolic model to assign the gene expression to the corresponding enzymes. The data was used to compute the fold changes between conditions and to classify the enzymes into up- or down-regulated using as threshold 1.3. Therefore, enzymes with a fold change above 1.3 are considered up-regulated and enzymes with a fold change below 1/1.3 are considered down-regulated.

We then developed an extended version of the method REMI (5), which integrates transcriptomics data into genome scale models for two conditions and analyzes the consistency of the data with the reaction rates in the network. The main assumption underlying REMI is that deregulations in the gene expression translate to deregulations of the corresponding enzyme abundance and therefore the reaction rate. With this, REMI imposes constraints so that if the gene associated to a reaction is upregulated, then its reaction rate must be higher. On the contrary, if the gene is downregulated the reaction rate will be lower. Then the method maximizes the number of reaction rates that can be simultaneously constrained in the network according to the fold change of the corresponding gene expressions between the two conditions [see (5) for further details]. In this work, we extended the REMI method to be able to integrate data for four conditions at the same time. Therefore, we generated four models of the redTCELL network, one per treatment, and we integrated the six ratios corresponding to the pairwise comparison of the four conditions (Table 2). Based on the data there are a total of 5668 reactions deregulated (up or down) when we perform pairwise comparisons between the treatments. Out of them, our method predicts that 5367 reaction rates can be simultaneously constrained in the network to satisfy the gene expression ratios (Table 2), integrating over 93% of all the deregulations between treatments.

| Fold change of comparisons between treatments | Theoretical number of deregulated reactions based on the expression data | Reactions in the model with rates consistent with the data |
|---|---|---|
| IL-2$^{50U}$ vs IL-15$^{50U}$ | 1096 | 1029 |
| IL-2v$^{5U}$ vs IL-15$^{50U}$ | 1108 | 1062 |
| IL-2v$^{50U}$ vs IL-15$^{50U}$ | 1168 | 1108 |
| IL-2v$^{5U}$ vs IL-2$^{50U}$ | 992 | 933 |
| IL-2v$^{50U}$ vs IL-2$^{50U}$ | 579 | 553 |
| IL-2v$^{50U}$ vs IL-2v$^{5U}$ | 725 | 682 |

***Table 2. Statistics of metabolic reaction deregulations**. Number of deregulated reactions based on the data and consistency after integration in the model.*

### Metabolic reaction fluxes representative of each CD8$^+$ T cell treatment

We fixed the ratios of reactions consistent with the network, and we use the Artificial Centering Hit-and-Run sampler (ACHR) to sample 100K points from the solution space. We then computed for each reaction the mean and the mean fold change of the populations of samples between two conditions and we use it as representative fluxes for the analysis.

#### Extraction of minimal networks for metabolic task enrichment analysis

Seeking to study the proliferative phenotype of T cells using the redTCELL metabolic model, we defined five metabolic tasks associated to growth, i.e., DNA synthesis, RNA synthesis, protein synthesis, lipid synthesis and energy production.

Using the composition of the biomass reaction in redTCELL, we extracted the four parts representing the synthesis of the mentioned macromolecules, and we considered the ATPS reaction of the electron transport chain for the synthesis of ATP. Using MiNEA, a previously developed approach (6)[cite MINEA Pandey et al. *PLOS Comp Biol* 2019], we generated minimal networks for the five metabolic tasks. To this end, we formulated a mixed integer linear program (MILP) and we identified the minimum number of reactions required to synthesize each macromolecule and possible alternatives.

***Table 3. Minimal networks****. Size and number of alternatives of minimal networks generated for the five tasks associated to the proliferative phenotype. The minimal size is the minimum number of reactions that are required to synthesize each molecule and the number of alternatives represents different biochemical pathways that cells can use for this synthesis.*

| Metabolic Task | Minimal network size | Number of alternatives (Smin+1) |
|---|---|---|
| ATP production (ATPS) | 25 | 165 |
| DNA synthesis | 113 | 268 |
| RNA synthesis | 102 | 183 |
| Protein synthesis | 43 | 1000 |
| Lipid synthesis | 117 | 14 |

Next, we performed minimal network enrichment analysis using either the gene expression data or the representative of the metabolic fluxes computed by sampling after integrating the transcriptomics data into the redTCELL model.

#### Downstream signaling network for IL-2 receptor

We used REACTOME (https://reactome.org/) (7) to identify 29 signaling pathways downstream the IL-2 receptor and 16 pathways related to cell cycle.

We mapped the phospho-proteomics data to the species of the selected pathways, and we performed enrichment analysis by using the fold change abundance of the phosphorylated molecules in the T cells stimulated vs unstimulated.

#### Metabolomics analysis

$CD8^+$OT1 T cells were *in vitro* expanded in presence of IL-2v (5 and 50 IU/mL), IL-2 (50 IU/mL) or IL-15 (50 IU/mL) for 10 days. Supernatants and cells were collected, and a multiple pathway targeted metabolomics analysis was performed by the Metabolomics Unit at the University of Lausanne, Switzerland.

For metabolite extraction, supernatants derived from the cell cultures were extracted by the addition of ice-cold MeOH. Extracts were centrifuged for 15 min at 15,000 rpm at 4°C and the resulting supernatant was collected and transferred into a LC-MS vial for analysis. Also, $CD8^+$OT1 cells were pre-extracted and homogenized by the addition of MeOH: H2O (4:1) in the Cryolys Precellys 24 sample Homogenizer (2 x 20 sec at 10,000 rpm, Bertin Technologies) with ceramic beads. The bead beater was air-cooled down at a flow rate of 110 L/min at 6 bar. Homogenized extracts were centrifuged for 15 min at 4,000 g at 4°C. The resulting supernatant was collected and evaporated to dryness in a vacuum concentrator (LabConco). Dried sample extracts were resuspended in MeOH: H2O (4:1, v/v) according to the total protein content. The protein pellets were evaporated and lysed in 20 mM Tris-HCl (pH 7.5), guanidine hydrochloride (4 M), NaCl (150 mM), $Na_2EDTA$ (1 mM), EGTA (1 mM), 1% Triton, sodium pyrophosphate (2.5 mM), β-glycerophosphate (1 mM), $Na_3VO_4$ (1 mM) and leupeptin (1 µg/mL) using the Cryolys Precellys 24 sample Homogenizer (2 x 20 seconds at 10,000 rpm, Bertin Technologies) with ceramic beads. BCA Protein Assay Kit (Thermo Fisher Scientific) was used to determine (A562 nm) total protein concentration (Hidex).

Extracted cell and supernatant samples were analyzed by hydrophilic interaction liquid chromatography coupled to tandem mass spectrometry (HILIC-MS/MS) (8,9) in both positive and negative ionization modes by using a 6495 triple quadrupole system (QqQ) interfaced with 1290 UHPLC system (Agilent Technologies). The chromatographic separation in positive mode was performed in an Acquity BEH Amide, 1.7 µm, 100 mm x 2.1 mm inner diameter column (Waters). Mobile phase consisted of A (20 mM ammonium formate and 0.1 % formic acid in water) and B (0.1 % formic acid in acetonitrile). The linear gradient elution from 95% B (0-1.5 min) down to 45% B was applied (1.5-17 min) and these conditions were held for 2 min. Then, the column was re-equilibrated for 5 min at the initial chromatographic conditions. The flow rate was 400 µl/min, column temperature 25°C and sample injection volume 2 µl. In negative mode, a SeQuant ZIC-pHILIC, 100 mm, 2.1 mm inner diameter and 5 µm particle size column (Merck) was used. In this case, the mobile phase was consisted of A (20 mM ammonium acetate and 20 mM NH4OH in water, at pH 9.7) and B (100% acetonitrile). The linear gradient elution from 90% (0-1.5 min) to 50% B (8-11 min) down to 45% B (12-15 min). Finally, the column was re-equilibrated for 9 min at the initial chromatographic conditions (10). The flow rate was 300 µl/min, column temperature 30°C and sample injection volume 2µl. In both analyses, electrospray ionization source conditions were set as follows: dry gas temperature 290 °C, nebulizer 35 psi and flow 14 L/min, sheath gas temperature 350°C and flow 12 L/min, nozzle voltage 0 V, and capillary voltage ±2,000 V. As acquisition mode Dynamic Multiple Reaction Monitoring (DMRM) was used, with a total cycle time of 600 ms. Optimized collision energies for each metabolite were applied.

Pooled quality control samples were analyzed periodically throughout the overall analytical run to assess the quality of the data, correct the signal intensity drift and remove the peaks with poor reproducibility (coefficient of variation > 30%) (10). In addition, a series of quality controls diluted with methanol (100%, 50%, 25%, 12.5% and 6.25%) were prepared. Then, metabolites were selected also considering the linear response on the diluted quality control series.

Raw LC-MS/MS data were processed using the Agilent Quantitative analysis software (v.B.07.00, MassHunter Agilent technologies). Relative quantification of metabolites was based on extracted ion chromatogram areas for the monitored MRM transitions. The obtained tables (with peak areas of detected metabolites) were exported to R software (http://cran.r-project.org/) and signal intensity drift correction and noise filtering was done within the MRMPROBS software.

**Protein phosphorylation analysis.**

Naïve CD8$^+$OT1 cells were stimulated *in vitro* on day 0 with anti-CD3/anti-CD28 coated beads and expanded up to 8 days in presence of IL-15 (10 ng/mL) to maximize the yield of $T_{CM}$-like cells. For studying the protein phosphorylation, an untargeted approach based on phospho-proteomics by mass spectrometry was performed by the Protein Analysis Facility at the University of Lausanne, Switzerland (CIG, UNIL). Briefly, on day 8, the cells were FBS starved for 2 hours and subsequently stimulated for 20 min with IL-2, IL-2v or IL-15 at 5 and

50 IU/mL. Next, the cells were lysed, digested, and the resultant peptide samples were desalted using Oasis HLB plates (Waters). The phospho-peptide enrichment was carried out using a titanium dioxide ($TiO_2$) standard protocol. Analysis based on phospho-site abundance relative to unstimulated OT1 cells was performed.

In addition, a targeted approach by phospho-flow cytometry was carried out by staining with the following antibodies: p-STAT5 (T694), p44/42 MAPK, p-AKT(T308), p-AKT (S473), p-S6 (S235/236), pFoxO1 (T24) / FoxO3a (T32), p-GSK-3β (S9) from Cell Signaling Technologies. Previously *in vitro* expanded and FBS starved $CD8^+$OT1 cells were stimulated with the mentioned cytokines at 5, 10, 25 and 50 IU/mL for 20 min. Also, in indicated experiments, $CD8^+$OT1 cells were stimulated with IL-2v (5 IU/mL) or IL-15 (50 IU/mL) for the indicated times. Besides, phosphorylation of mentioned proteins was measured in human $CD8^+$ T cells. Cells were activated and expanded *in vitro* in presence of IL-15 (10 ng/mL) for 8 days, and then, FBS starved for 2 hours and stimulated with IL-2, IL-2v or IL-15 (5 and 50 IU/mL) for 20 min. Data were obtained with an LSRII flow cytometer (BD Biosciences).

**Immunoblotting**

$CD8^+$ OT1 T cells were activated and expanded in T-cell medium supplemented with IL-2 (50 IU/mL) for 6 days. The cells were subsequently FBS and cytokine-starved for 2 hours and then cultured in presence of IL-2 (0.5, 5, 50 and 500 IU/mL) or IL-2v (5 and 50 IU/mL). The cells were collected after 15 minutes, lysed with RIPA buffer supplemented with Halt phosphate/protease inhibitors (Thermo Fisher Scientific), and boiled at 95°C for 10 minutes with Bolt LDS sample buffer and reducing agent (Thermo Fisher Scientific). Protein samples were separated by SDS-PAGE and transferred to PVDF membranes using the iBlot2 system (Thermo Fisher Scientific). Membranes were incubated with the following antibodies according to the manufacturer's instructions: p-JAK1 (74129), JAK1 (3344), p-JAK3 (5031), JAK3 (8863), obtained from Cell Signaling Technology, and β-actin (sc-47778) from Santa Cruz. Bound antibodies were detected with horseradish peroxidase (HRP)-conjugated secondary antibodies. Proteins were visualized using chemiluminescent substrate (ECL, GE healthcare). Some membranes were stripped with stripping buffer (Tris 62.5mM pH6.7, SDS 2%, 0.1M β-mercaptoethanol), for 30 minutes at 50°C. Phosphorylation status was calculated by dividing the signal of the phosphorylated protein by the signal of the matched total protein. Protein levels were quantified using the ImageJ software.

**Tumor inoculation and Adoptive T-cell transfer**

C57BL/6 mice were injected subcutaneously with $1x10^5$ B16.OVA tumor cells. When tumors reached an average volume between 50-70 $mm^3$, mice received 4 Gy of sub-lethal total body irradiation (TBI) and were grouped (n ≥ 8 mice/group). On days 10 and 14, mice were treated with *i.v* transfer of $2.5x10^6$ $CD8^+CD45.1^+$ OT1 cells cultured with IL-2v (5 IU/mL) or IL-2 (50 IU/mL). Tumors and tumor-draining lymph node were collected on days 2, 6 and 10 after ACT for subsequent analyses. For exploring the engraftment of transferred T cells, C57BL/6 $CD45.2^+$ mice were irradiated and treated with a total of $2.5x10^6$ $CD8^+CD45.1^+$ OT1 cells previously *in vitro* expanded in presence of IL-2v (5 IU/mL) or IL-2 (50 IU/mL). After 24 hours of ACT, exogenous $CD45.1^+$ OT1 cells were isolated from spleens and analyzed by flow cytometry. Additionally, in indicated experiments, the ACT consisted of $CD8^+CD45.1^+$OT1 cells engineered to secrete IL-2v or IL-15.

In experiments evaluating the impact on T-cell performance of IL-15 and IL-7 combined with IL-2v (5 IU/mL), B16-OVA tumor-bearing CD56Bl/6 mice were adoptively transferred with

$CD8^+CD45.1^+$ cells previously *in vitro* expanded with IL-2v (5 IU/mL) with or without IL-15/IL-7 (10 ng/mL) or IL-2 (50 IU/mL) plus IL-15/IL-7.

Mice were carefully monitored, and tumor length (L; greatest longitudinal measurement) and width (W; greatest transverse measurement) were measured by caliper every 2-3 days. Tumor volumes (V) were calculated using the formula: $V = (L \times W^2)/2$. Mice were sacrificed once tumors reached 1000 $mm^3$, or according to regulations if they became distressed or moribund, or at described time-points post-ACT.

**Statistical analyses**

All analyses were performed using GraphPad Prism version 7.03 (Graph Pad Software). Data are expressed as mean ± standard deviation (SD) or mean ±standard error of the mean (SEM), as indicated. For the analysis of normality of the experimental groups, D´Agostino and Pearson Omnibus tests were used. For the analysis of homoscedasticity, the Bartlett´s test for equal variances was used. Statistical analysis of two experimental groups was assessed using a parametric unpaired 2-tailed Student's t test or unpaired 2-tailed Student's t test with Welch's correction, as appropriate. Multiple comparisons were assessed by using One-Way Analysis of Variance (ANOVA) followed by Tukey post-test or non-parametric Kruskal-Wallis test and Dunn's multiple comparison post-test. Analysis of survival was performed using a Log-rank Mantel-Cox test. In all analysis, $p$ values $< 0.05$ were considered significant (*: $p < 0.05$; **: $p < 0.01$; ***: $p < 0.001$).

Low-dose IL-2v couples CD8+T cell expansion to proliferation while preserving mitochondrial fitness via tonic IL-2R signaling and co-activation of memory/stemness and proliferation transcriptional programs.

IL-2v when applied at low-doses couples CD8+T cells expansion to proliferation while preserving mitochondrial fitness via low-intensity tonic IL-2R signaling and co-activation of memory/stemness and proliferation transcriptional programs

Reference:

1. Gillis, S., Ferm, M. M., Ou, W., and Smith, K. A. (1978) T-Cell Growth-Factor - Parameters of Production and a Quantitative Microassay for Activity. *Journal of Immunology* **120**, 2027-2032
2. Lanitis, E., Rota, G., Kosti, P., Ronet, C., Spill, A., Seijo, B., Romero, P., Dangaj, D., Coukos, G., and Irving, M. (2021) Optimized gene engineering of murine CAR-T cells reveals the beneficial effects of IL-15 coexpression. *J Exp Med* **218**
3. Masid, M., Ataman, M., and Hatzimanikatis, V. (2020) Analysis of human metabolism by reducing the complexity of the genome-scale models using redHUMAN. *Nat Commun* **11**, 2821
4. Brunk, E., Sahoo, S., Zielinski, D. C., Altunkaya, A., Drager, A., Mih, N., Gatto, F., Nilsson, A., Preciat Gonzalez, G. A., Aurich, M. K., Prlic, A., Sastry, A., Danielsdottir, A. D., Heinken, A., Noronha, A., Rose, P. W., Burley, S. K., Fleming, R. M. T., Nielsen, J., Thiele, I., and Palsson, B. O. (2018) Recon3D enables a three-dimensional view of gene variation in human metabolism. *Nat Biotechnol* **36**, 272-281
5. Pandey, V., Hadadi, N., and Hatzimanikatis, V. (2019) Enhanced flux prediction by integrating relative expression and relative metabolite abundance into thermodynamically consistent metabolic models. *PLoS Comput Biol* **15**, e1007036
6. Pandey, V., and Hatzimanikatis, V. (2019) Investigating the deregulation of metabolic tasks via Minimum Network Enrichment Analysis (MiNEA) as applied to nonalcoholic fatty liver disease using mouse and human omics data. *PLoS Comput Biol* **15**, e1006760
7. Gillespie, M., Jassal, B., Stephan, R., Milacic, M., Rothfels, K., Senff-Ribeiro, A., Griss, J., Sevilla, C., Matthews, L., Gong, C., Deng, C., Varusai, T., Ragueneau, E., Haider, Y., May, B., Shamovsky, V., Weiser, J., Brunson, T., Sanati, N., Beckman, L., Shao, X., Fabregat, A., Sidiropoulos, K., Murillo, J., Viteri, G., Cook, J., Shorser, S., Bader, G., Demir, E., Sander, C., Haw, R., Wu, G., Stein, L., Hermjakob, H., and D'Eustachio, P. (2022) The reactome pathway knowledgebase 2022. *Nucleic Acids Res* **50**, D687-D692
8. Gallart-Ayala, H., Konz, I., Mehl, F., Teav, T., Oikonomidi, A., Peyratout, G., van der Velpen, V., Popp, J., and Ivanisevic, J. (2018) A global HILIC-MS approach to measure polar human cerebrospinal fluid metabolome: Exploring gender-associated variation in a cohort of elderly cognitively healthy subjects. *Anal Chim Acta* **1037**, 327-337
9. Medina, J., van der Velpen, V., Teav, T., Guitton, Y., Gallart-Ayala, H., and Ivanisevic, J. (2020) Single-Step Extraction Coupled with Targeted HILIC-MS/MS Approach for Comprehensive Analysis of Human Plasma Lipidome and Polar Metabolome. *Metabolites* **10**
10. Broadhurst, D., Goodacre, R., Reinke, S. N., Kuligowski, J., Wilson, I. D., Lewis, M. R., and Dunn, W. B. (2018) Guidelines and considerations for the use of system suitability and quality control samples in mass spectrometry assays applied in untargeted clinical metabolomic studies. *Metabolomics* **14**, 72
